# Simple, Functional, Inexpensive Cell Extract for *in vitro* Prototyping of Proteins with Disulfide Bonds

**DOI:** 10.1101/2019.12.19.883413

**Authors:** Jared L. Dopp, Nigel F. Reuel

**Affiliations:** Iowa State University Chemical and Biological Engineering

**Keywords:** Disulfide bond, cell-free protein synthesis, luciferase, chitinase, T7 SHuffle

## Abstract

*In vitro* expression of proteins from *E. coli* extract is a useful method for prototyping and production of cytotoxic or unnatural products. However, proteins that have multiple disulfide bonds require custom extract that, to date, requires careful addition of exogenous isomerase enzymes or the use of expensive commercial kits. This cost and complexity currently limit access to some groups who wish to rapidly prototype proteins with disulfide bonds. Herein, we present a simple solution that does not require addition of supplemental enzymes. We use a commercially available SHuffle T7 Express *lysY* strain of *E. coli* that expresses both T7 RNAP and DsbC isomerase enzymes. We experimentally determine optimal growth conditions (IPTG induction and harvest times) to balance overall productivity and efficiency of disulfide bond formation using a luciferase (from *Gaussia princeps*) that contains five disulfide bonds as our reporter protein. We also demonstrate the ability for rapid prototyping by screening the activity of four luciferase candidates against ten luciferin analogues. To display the broad applicability of the extract, three other enzymes containing ≥3 disulfide bonds (hevamine, endochitinase A, and periplasmic AppA) were also expressed from minimal genetic templates that had undergone rolling circle amplification and confirmed via activity assays.

## Introduction

Cell free protein synthesis (CFPS) has seen many recent applications in prototyping gene circuits (Green et al., 2017; Siegal-Gaskins et al., 2014; Sun et al., 2014), sensors (Mcnerney et al., 2019; Pardee et al., 2016a; Salehi et al., 2018; Soltani et al., 2018; Takahashi et al., 2018), and therapies (Cai et al., 2015; Pardee et al., 2016b; Salehi et al., 2016). Cell extract derived from *E. coli* is the most widely used with simple, efficient extract protocols, multiple supplement recipes, and clear experimental protocols available in literature (Dopp et al., 2019b; Dopp et al., 2019a; Dopp and Reuel, 2018; Kwon and Jewett, 2015; Levine et al., 2019; Sun et al., 2013; Yang et al., 2012). One major limitation of *E. coli* extract for protein expression is the inability to perform post-translational modifications. There has been much work to engineer glycosylation (Jaroentomeechai et al., 2018; Schoborg et al., 2018) and disulfide bond-formation (Kim and Swartz, 2004; Matsuda et al., 2013) pathways. However, these improvements increase the complexity of extract preparation, requiring careful fermentation and skilled augmentation with purified chaperones and enzymes that are expressed in other cells. Commercial kits containing supplements to improve disulfide bond formation are now available but significantly increase the cost per reaction (GeneFrontier; NEB). It is the central goal of this work to create a simple, single cell growth method for creating inexpensive, functional extract for rapid prototyping of proteins with multiple disulfide bonds. Such simple, inexpensive approaches have been shown to have a significant impact on the accessibility and rate of scientific innovation (Whitesides, 2010; Whitesides, 2013).

Many extracellular proteins of interest contain multiple disulfide bonds (Wong et al., 2011), such as therapeutic proteins like human growth hormone (Junnila and Kopchick, 2013), proinsulin (Haataja et al., 2016), antibodies (Saeed et al., 2017), and interleukin-6 (Snouwaert et al., 1991). Disulfide bonds are important because they can improve resistance to degradation by proteases and improve thermodynamic stability (Fass, 2012). Industrial enzyme producers and the biopharmaceutical industry use this strategy to engineer and improve wild type proteins (Lobstein et al., 2012; Matsuda et al., 2013). In eukaryotes, disulfide bonds are typically formed in the endoplasmic reticulum (ER) while folding in the periplasm of *E. coli* (Inaba, 2010; Patil et al., 2015). This typically leads to low yields (Ke and Berkmen, 2014; Mamathambika and Bardwell, 2008) and can result in proteins being sequestered to inclusion bodies that require refolding (Baneyx and Mujacic, 2004; Burgess, 2009; Middelberg, 2002). Since the periplasm is compromised during cell lysis, there is no separate compartment for these oxidative processes. Figure 1 depicts the oxidation, reduction, and isomerization processes undergone by thiol groups cysteine residues.

**Figure 1.**
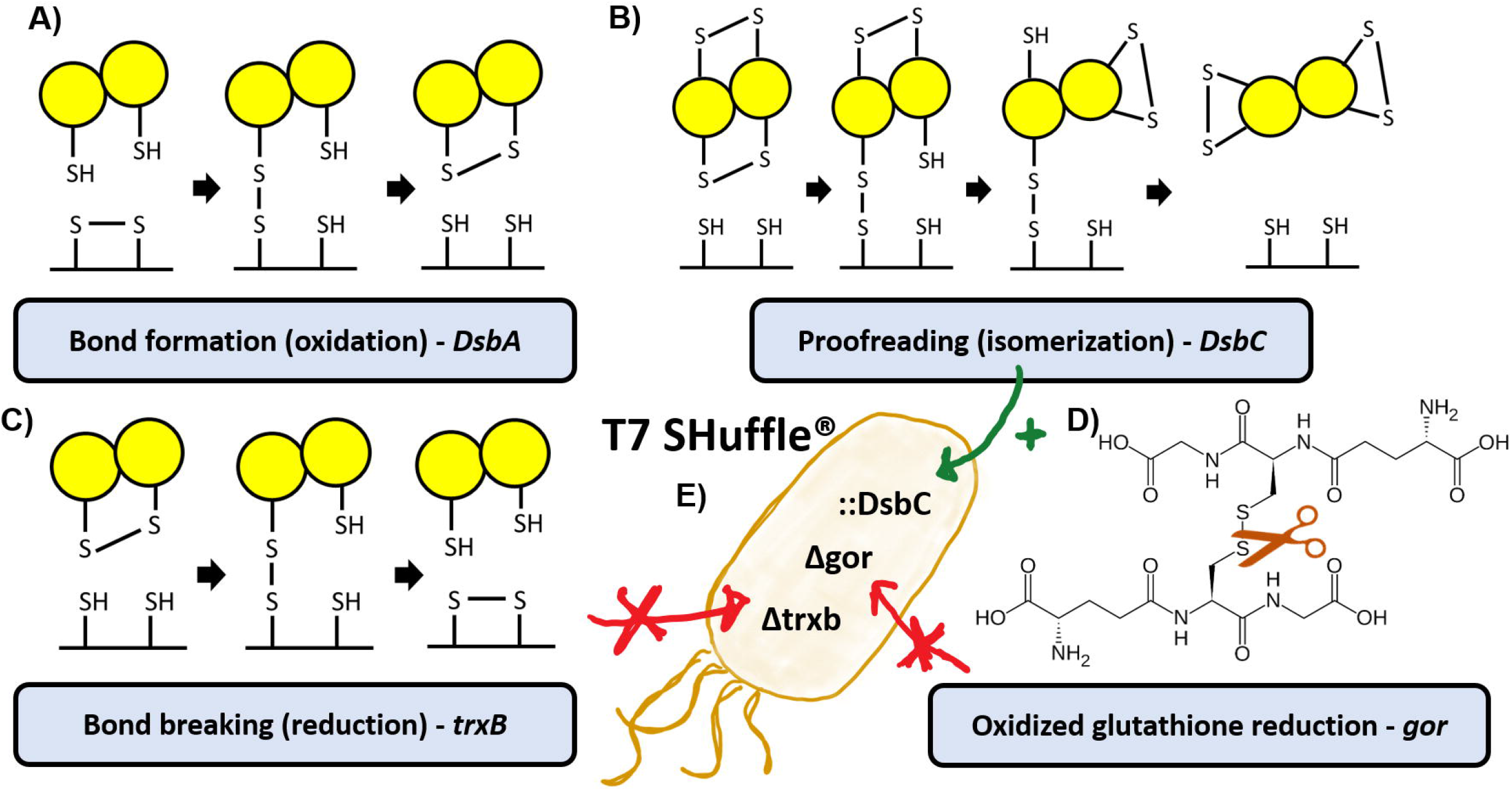
Schematic of mechanisms that affect disulfide bond formation and implications in the T7 Shuffle strain. (A) Oxidation via DsbA forms bonds between thiol groups on cysteines. (B) DsbC enzymes proofread proteins and isomerize disulfide bonds. (C) Reduction can occur via trxB and (D) gor enzymes that cleave disulfide bonds on the protein. (E) The T7 SHuffle^®^ strain is engineered to support disulfide bond formation by eliminating reducing enzymes and overexpressing the DsbC chaperone. This figure has been modified from a published schematic on disulfide bond formation (Ke and Berkmen, 2014).

With these issues in mind, researchers have taken steps to improve cytoplasmic expression (*in vivo* and *in vitro*) of proteins with disulfide bonds. In 2003, Kim and Swartz successfully expressed the protease domain of murine urokinase (6 disulfide bonds) using extract from the *E. coli* A19 strain (K12 derivative). This was achieved by manipulating the redox environment and adding disulfide bond isomerase C (DsbC) (Kim and Swartz, 2004). Soon thereafter, a truncated version of tissue type plasminogen activator (9 disulfide bonds) was successfully expressed. Decreasing the reaction temperature to 30 °C and adding the chaperone Skp improved expression (Yin and Swartz, 2004). Around the same time, the KC1 strain was developed to improve amino acid stability (arginine, tryptophan, and serine) during CFPS reactions (Michel-Reydellet et al., 2004). In 2005, an active GMCSF-sfFv fusion protein (4 disulfide bonds) was produced using the NMR2 strain, another modification from A19 (Michel-Reydellet et al., 2004; Voloshin and Swartz, 2005). More progress was made when KC1 was modified to create the KC6 mutant with enhanced cysteine stability due to the deletion of glutamate-cysteine ligase (*gshA*). This stabilization led to improved overall production but a massive increase in active murine urokinase and GM-CSF (Calhoun and Swartz, 2006). In order to support disulfide bond formation during CFPS using glucose as an energy source, the KGK10 strain was created by deleting the glutathione reductase (*gor*) gene from the KC6 strain. A hemagglutinin purification tag was also added to the C-terminus thioredoxin reductase gene (*trxB*) but its removal proved to be unnecessary. Murine urokinase and GM-CSF were produced but with lower yields due to glucose replacing phosphoenolpyruvate (PEP) as an energy source (Knapp et al., 2007). A direct comparison of KGK10 and KC6 using the PANOx-SP system showed few decreases in production but many improvements. It was also shown to be effective at expressing multiple scFv vaccine candidates showcasing the potential for CFPS in the therapeutic space (Goerke and Swartz, 2007). Around this same time, the Buchner group was able to express a full-length IgG antibody using CFPS. 507 ng/mL of active MAK33 was produced using two commercial kits (RTS100 HY and RTS500 HY) upon the addition of DsbC, GSSG, and GSH (Frey et al., 2008). In 2012, a correctly folded mutant of the flu virus hemagglutinin stem domain was produced using the KC6 strain. The number of disulfide bonds in the monomers of the stem domain was reduced from 4 to 2. In order to stabilize the trimer that forms the stem domain, cysteines were added to form disulfide bonds between monomers (Lu et al., 2014). This was an excellent display of using CFPS for protein engineering to improve protein production and stability by manipulating disulfide bond formation.

These advancements have led to industrial-scale applications of CFPS for producing therapeutics with disulfide bonds. Sutro Biopharma achieved a large milestone by demonstrating the successful scale-up of CFPS by producing recombinant human GM-CSF, along with other therapeutic proteins. They found that protein production using a cell-free system scales linearly as the reaction volume is increased (Zawada et al., 2011). In 2012, Sutro hit another milestone by demonstrating they’re Open Cell-Free System (OCFS) could be used for high throughput antibody engineering. In the process, they were able to optimize conditions for trastuzumab IgG1 (16 disulfide bonds) expression (Yin et al., 2012). Sutro then engineered a strain to improve mAb production by incorporating 2 chromosomal copies of DsbC and a plasmid with 2 copies of FkpA (Groff et al., 2014). This work has led to the production of the SBJY001 strain which requires less processing during extract production with the ability to produce tumor seeking antibody drug conjugates (Yin et al., 2017). Unfortunately, these strains require defined media, fermentation, and dialysis, and are not commercially available (Liu et al., 2005; Zawada and Swartz, 2005) which has limited their widespread impact.

To quantify impact of cell extract methods, we have quantified their use in scientific citations. The most used strain in CFPS, BL21 (DE3) Star, is a commercially available strain capable of producing functional lysate with minimal processing steps and has been widely cited (Fig 2). BL21 (DE3) Star, and similar BL21 strains, have been used to express proteins containing disulfide bonds with varying success. For example, BL21 (DE3) RIPL-Star was used to express human and murine proteins, each containing 2 disulfide bonds, without pretreating the extract. It was also shown that creating a fusion with the GB1 protein at the N terminus increases expression (Michel and Wüthrich, 2012b; Michel and Wüthrich, 2012a). More recently, BL21 (DE3) Star has been used to successfully express correctly folded hydrophobins (4 disulfide bonds) upon the addition of DsbC but no IAM pretreatment (Siddiquee et al., 2020). However, these BL21 (DE3) Star strains do not have modifications aimed at improving disulfide bond formation. The only commercial systems available are kits that provide expensive supplements for disulfide bond formation. The PURE*frex*^®^ 2.1 system, made of purified, recombinantly expressed proteins, is one of these commercial platforms capable of reliably producing proteins with disulfide bonds (Murakami et al., 2019). This reliability comes at a steep price with a cost of roughly US $0.57/μL of reaction and inclusion of the disulfide bond supplement raises the price to US $0.70/μL of reaction and has thus limited impact to those research groups that can afford such kits (Fig 2). On the other hand, creating CFPS components in house can reduce cost to $0.02/μL of reaction (Gregorio et al., 2019; Levine et al., 2019). The complexity and access limitations to KC6, KGK10, and SBJY001 are likewise revealed when counting citations of books, peer reviewed articles, and preprints from Google Scholar (Fig 2).

**Figure 2.**
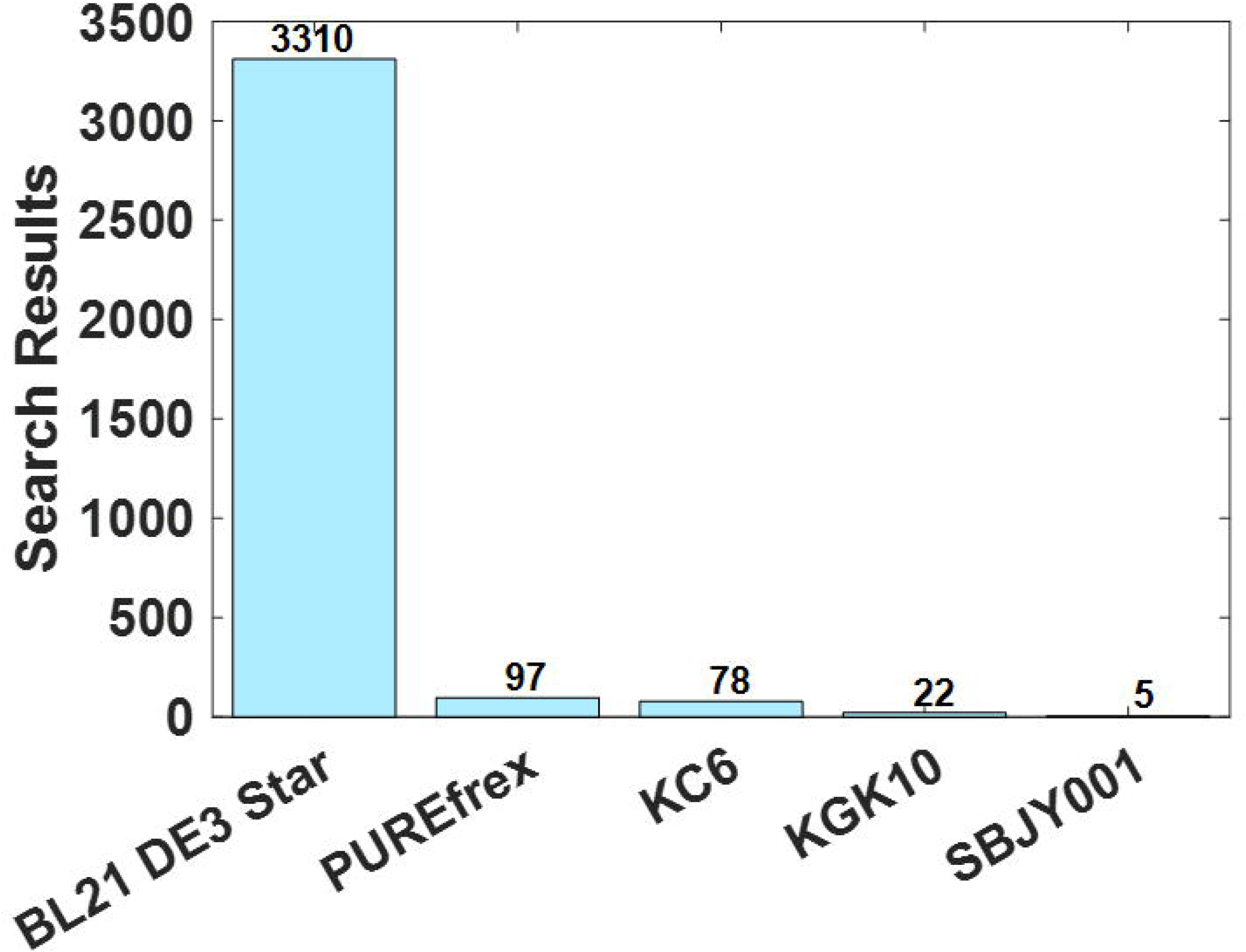
The number of search results from Google Scholar when searching cell free and the strain name between the years 2000 and 2020. The results from PUREfrex come from searching for PUREfrex alone since it is exclusively a cell-free platform. These results include mentions in published articles, preprints, and books. These numbers represent the results from a search performed February 21, 2020.

In line with our goal to make CFPS a simple, functional, inexpensive method that is readily transferable to any research group (Dopp et al., 2019b; Dopp et al., 2019a; Dopp and Reuel, 2018), we desired to identify and optimize a strain that could be used to produce extract capable of forming disulfide bonds without the need for supplementation with exogenous enzymes. Ideally, the strain should be grown in a simple shake flask and would not require a run-off reaction or dialysis. We identified a commercially available *E. coli* strain that has been optimized to support *in vivo* disulfide bonded protein expression in the cytoplasm, T7 SHuffle^®^ by New England BioLabs (NEB). The SHuffle strain is derived from ER2566 and has been modified to constitutively express disulfide bond isomerase C (DsbC) in the cytoplasm and overexpress T7 RNA polymerase (T7 RNAP) upon induction by isopropyl β-d-1-thiogalactopyranoside (IPTG) (Fig 1). The SHuffle strain also eliminates unwanted reduction pathways in the cytoplasm due to deletion of the *trxB* and *gor* genes (Fig 1E) (Anton et al., 2016; Lobstein et al., 2012; Robinson et al., 2015).

To understand the significance of these modifications, it is important to have an understanding of the intracellular redox environment. The indiscriminate oxidation of cysteines is carried out by disulfide bond isomerase A (DsbA) which forms disulfide bonds between successive cysteines (Fig 1A). DsbC allows for thermodynamically driven proof-reading by isomerizing disulfide bonds that have been formed between sub-optimal cysteines, thus allowing for continued folding to the proper state and disulfide linking (Fig 1B). In an unmodified strain, thioredoxin reductase will reduce disulfide bonds or prevent them from forming. Also, in unmodified strains, glutathione reductase will reduce oxidized glutathione (Fig 1D), thus removing its ability to oxidize thiol groups on cysteines (Ke and Berkmen, 2014; Lobstein et al., 2012).

In this work, we investigate the potential of T7 SHuffle^®^ as a platform for CFPS based prototyping of proteins with multiple disulfide bonds. We optimize the extract preparation based on our scalable techniques previously used for BL21 (DE3) Star cells (Dopp and Reuel, 2018). Luciferase from *Gaussia princeps* (Gluc) is used as a disulfide bonded reporter protein for extract optimization due to its efficient luminescence and previous application in CFPS (Goerke et al., 2008; Liu et al., 2014; Yu et al., 2018). We show that T7 SHuffle^®^ is superior to BL21 (DE3) Star when expressing Gluc (5 disulfide bonds). We then demonstrate the broader applicability of this extract by screening 4 luciferase candidates with >90% homology to Gluc against a panel of 10 luciferase substrates. We further demonstrate prototyping with this extract by expressing three additional enzymes with disulfide bonds: hevamine from the rubber tree *Hevea brasiliensis* (Bokma et al., 2002), endochitinase A (ChitA) from *Zea mays* (Hawkins et al., 2015), and the periplasmic acid phosphatase, phytase (AppA) from *Escherichia coli* (Berkmen et al., 2005) which have 3, 7, and 4 disulfide bonds respectively. Some practical applications of the two chitinases include agricultural pathogen control, contribution to asthma, and formation of chitooligosaccharides (Hamid et al., 2013). Moreover, we introduce a custom fusion protein that allows us to measure the ratio between oxidative potential and general productivity of an extract.

## Materials and Methods

### Extract Preparation

#### T7 SHuffle^®^

Extract from the SHuffle^®^ T7 Express *lysY* strain (gift from the Mansell lab) was prepared according to previously established protocols (Dopp and Reuel, 2018). Cells were grown and harvested in 200 mL 2x YTPG media using a 500 mL shake flask at 37°C and 230 RMP. Cultures were harvested at different growth/induction times according to the DoE design (Supplementary Table 1). The cells were centrifuged for 15 minutes at 5,000 xg and 4°C. The cell pellets were collected and washed in 10 mL of cold buffer A and centrifuged for 15 minutes at 4,255 xg and 4°C (Dopp et al., 2019a). The supernatant was discarded, and the wet cell mass was measured before freezing the cells at −80°C. Cells were thawed and resuspended in 1 mL of buffer A per gram of wet cell mass. Suspended cells were tip sonicated (QSonica) in 1 mL aliquots at −4°C using a 10 sec on/off pulse and 50% amplitude until the energy input reached 532 J (Kwon and Jewett, 2015). Sonicated cell suspensions were then centrifuged for 10 minutes at 4°C and 12,000 xg to create S12 extract. The supernatant (S12 extract) was collected and stored at −80°C in 45 uL aliquots. Once the optimum growth/induction times were determined for 200 mL cultures, Matlab was used to fit 200 mL and 1 L culture growth curves. The regression used was the Verhulst-Pearl equation (VPE) used in other similar studies (Supplementary Figure 1) (Wilding et al., 2018). These curve fits were used to correlate the optimum at 200 mL with a predicted optimum at 1 L. Optimized 1 L cultures were then grown in 2.5 L Tunair flasks and harvested according to the aforementioned protocol but were lysed using a French press homogenizer (Avestin EmulsiFlex C3) according to previous protocols (Dopp and Reuel, 2018). Frozen cells were resuspended in 1 mL buffer A + 1 mL ddH2O per 1 g of wet cell mass. This double dilution reduces the viscosity and maintains buffer composition upon reconstitution. The suspension was then fed through the French press and lysed at 20,000 - 30,000 psig. The crude lysate was then centrifuged for 10 min at 12,000 xg and 4°C. The supernatant was then collected, frozen at −80°C, and lyophilized in a VirTis pilot lyophilizer (SP Scientific) overnight. The resulting powder was collected, weighed, and stored at −80°C. Concentrated extract was obtained by re-lyophilizing extract and adding less water. Dilute extract was obtained by adding more water.

#### KGK10

The KGK10 strain (gift from Swartz lab) was grown in a 500 mL shake flask and monitored every 15 min to obtain a growth curve. This growth curve was used to find the point at which the rate of growth was the highest. KGK10 was then grown and harvested at 240 min (Supplementary Figure 2). The cells were centrifuged for 15 minutes at 5,000 xg and 4°C. The cell pellets were collected and washed in 10 mL of buffer A and centrifuged for 15 minutes at 4,255 xg and 4°C. The supernatant was discarded, and the wet cell mass was measured before freezing the cells at −80°C. Cells were thawed and resuspended in 1 mL buffer A + 1 mL ddH_2_O per 1 g of wet cell mass, identical to the method used for SHuffle extract. The suspension was then fed through the French press and lysed at 20,000 - 30,000 psig. The crude lysate was then centrifuged for 10 min at 12,000 xg and 4°C. The supernatant was then collected, frozen at - 80°C, and lyophilized in a VirTis pilot lyophilizer (SP Scientific) overnight. The resulting powder was collected, weighed, and stored at −80°C.

### DNA Amplification

All Gluc and sfGFP reporter protein expression experiments were carried out using plasmids. The sfGFP plasmid used was pJL1-sfGFP (gift from the Jewett lab) and the Gluc plasmid was pET24a-Gluc-6H (gift from the Swartz lab).

The production of minimal genetic templates for prototyping new enzymes has been described in previous work (Dopp et al., 2019b). In brief, the gene of interest was codon optimized using IDT’s codon optimization tool and purchased in the form of a gene fragment. The gene fragment was then amplified using OneTaq (NEB), digested to form sticky ends with HindIII (NEB), ligated with T4 ligase (NEB), and isothermally amplified with TempliPhi (GE Healthcare). This was all done in under 24 hr using a standard C1000 Touch Thermal Cycler (BioRad). The sequences for all linear and minimal genetic templates used are in the accompanying supplementary information.

### Cell-Free Reaction

#### T7 SHuffle^®^

The supplement recipe used for the designed experiments and initial tests is a modified version of the PANOx-SP system that is improved to form disulfide bonds (Dopp et al., 2018). Since the SHuffle strain can be induced to produce T7 RNAP and DsbC, these proteins were omitted from the reaction mix. The cell-free reaction included: 1.2 mM ATP, 0.85 mM each of GMP, UMP, and CMP, 30 mM phosphoenolpyruvate (Roche), 130 mM potassium glutamate, 10 mM ammonium glutamate, 12 mM magnesium glutamate, 1.5 mM spermidine, 1 mM putrescine, 34 μg/mL folinic acid, 171 μg/mL *E. coli* tRNA mixture (Roche), 4 mM oxidized glutathione (GSSG), 1 mM reduced glutathione (GSH), 2 mM each of 20 unlabeled amino acids, 0.33 mM NAD, 0.27 mM Coenzyme A (CoA), 4 mM potassium oxalate, 57 mM HEPES-KOH buffer (pH 7.5), 15 ng/uL plasmid, 0.24 volumes of E. coli S12 extract. Reactions were carried out in a 384 black-walled, flat-bottom well plate and shaken at 300 rpm using a ThermoMixer^®^ (Eppendorf). All proteins were expressed at 30°C unless otherwise stated. Gluc, hevamine, rChiA, and AppA reactions were run for 16 hrs, unless otherwise stated. The fluorescent proteins (sfGFP and rxYFP-mCherry) were expressed for 6 hrs. Reactions for the fluorescent proteins were carried out in a Synergy Neo2 HTS Multi-Mode Microplate Reader (BioTek) at 282 cpm.

#### KGK10

The cell-free reaction mixture for the KGK10 strain is identical to that used for the T7 SHuffle^®^ with the addition of 100 μg/mL DsbC (GeneFrontier) and 3.33 Units/uL T7 RNAP (Roche). Previously prepared KGK10 extract fermented and processed by the Swartz lab was tested alongside the in-house KGK10 extract prepared by shake flasks.

#### PURE*frex*^®^

The components for the PURE*frex* reactions were scaled down to suit 15 uL reactions. The components from the PURE*frex*^®^ 2.1 (GeneFrontier) were supplemented with DsbC and GSSG from the DS Supplement kit (GeneFrontier) according to manufacturer’s protocols.

## Assays

### Bioluminescence

The Gluc assay is based on previous work in the Swartz lab (Goerke et al., 2008). 3 μL of CFPS reaction was diluted in 100 μL nickel affinity EB buffer containing 1% w/v bovine serum albumin (BSA). The activity was measured by adding 2 μL of diluted sample to 100 μL assay buffer with 1 μL of 0.5 μg/μL coelenterazine (NanoLight Technology) dissolved in ethanol and immediately mixing before reading the luminosity. The sample buffer consisted of PBS pH 7.4 with 0.01% v/v Tween 20. Readings taken in a U-bottom 96 well plate. Luminescence was measured every 10 sec for 60 sec using a Synergy Neo2 HTS Multi-Mode Microplate Reader (BioTek). The plate reader settings used a 1536 filter, top optics position, and a gain of 100.

Coelenterazine analogues for the luciferase screening experiment were ordered from BioTium and suspended at a concentration of 0.5 μg/μL in the suggested solvent (ethanol or methanol). A water-soluble analogue was ordered from NanoLight Technology and dissolved at 0.5 μg/μL in ddH_2_O.

### Fluorescence

Fluorescence measurements were also taken using a Synergy Neo2 HTS Multi-Mode Microplate Reader (BioTek). Readings were taken every 5 min. The fluorescence of sfGFP was measured at an excitation of 485□nm and emission of 528□nm using a ±20 bandpass window and 61□gain setting. The reaction was also stirred in orbital motion at 237 cpm. For rxYFP, readings were taken at an excitation of 512 and emission of 523 using a ±5 bandpass window. Measurements for mCherry were taken at an excitation of 587 ±11 and an emission of 610 ±10.

### Enzyme Activity

Hevamine activity was determined by adding 10 μL CFPS reaction to 90 μL McIlvaine buffer pH 6.0 with 10 μL of 1 mM 4-methylumbelliferyl β-D-N,N′,N″-triacetylchitotrioside (4-MUF-TriNAG). rChiA activity was determined by adding 10 μL CFPS reaction to McIlvaine buffer pH 4.0 with 10 μL 4-methylumbelliferyl N-acetyl-β-D-glucosaminide (4-MUF-NAG). Fluorescence was measured at an excitation of 360 and emission of 445 with a bandpass window of ±20. Measurements were taken every 60 sec for 2 hrs in a black walled, flat bottom 384 well plate. AppA activity was determined by adding 10 μL CFPS reaction to 100 μL glycine buffer (250 mM glycine + 25 mM p-nitrophenyl phosphate) titrated to pH 3.0 with HCl. Absorbance readings were taken at 410 nm every 20 sec for 5 min in a 96 well plate at 37°C.

## Results and Discussion

### Designed Experiments

To optimize the growth conditions of the SHuffle strain, we performed two, face-centered-cubic (FCC) response surface design of experiments (DoE). This is an established statistical method to optimize an experimental outcome, with minimal experimental runs while accounting for cross-effects of input variables (Ferreira et al., 2007). In this case, the input variables under study are time of induction with ITPG to produce T7 RNAP and DsbC proteins as well as the overall time of harvest. The outcome to optimize is the amount of protein: in the first DoE, we measure the yield of sfGFP to compare to other strains (namely our work with BL21 DE3 Star cells); in the second, we measure the level of correctly folded Gluc to observe the level of disulfide-bonded protein yield. Gluc is a luciferase with five disulfide bonds and a strong luminescent signal making it a convenient reporter for extract optimization (Wu et al., 2015; Yu et al., 2018). In both cases, we constrained the growth times to encompass a single workday (< 7 hr growth time, as pelleting, washing, and storage must be completed that same day). Each response surface design yielded 13 runs, of which 5 were replicated center points to measure experimental drift. The measured values were then fit to quadratic models and the response surfaces are presented as contour plots (Fig 3 A-B).

**Figure 3.**
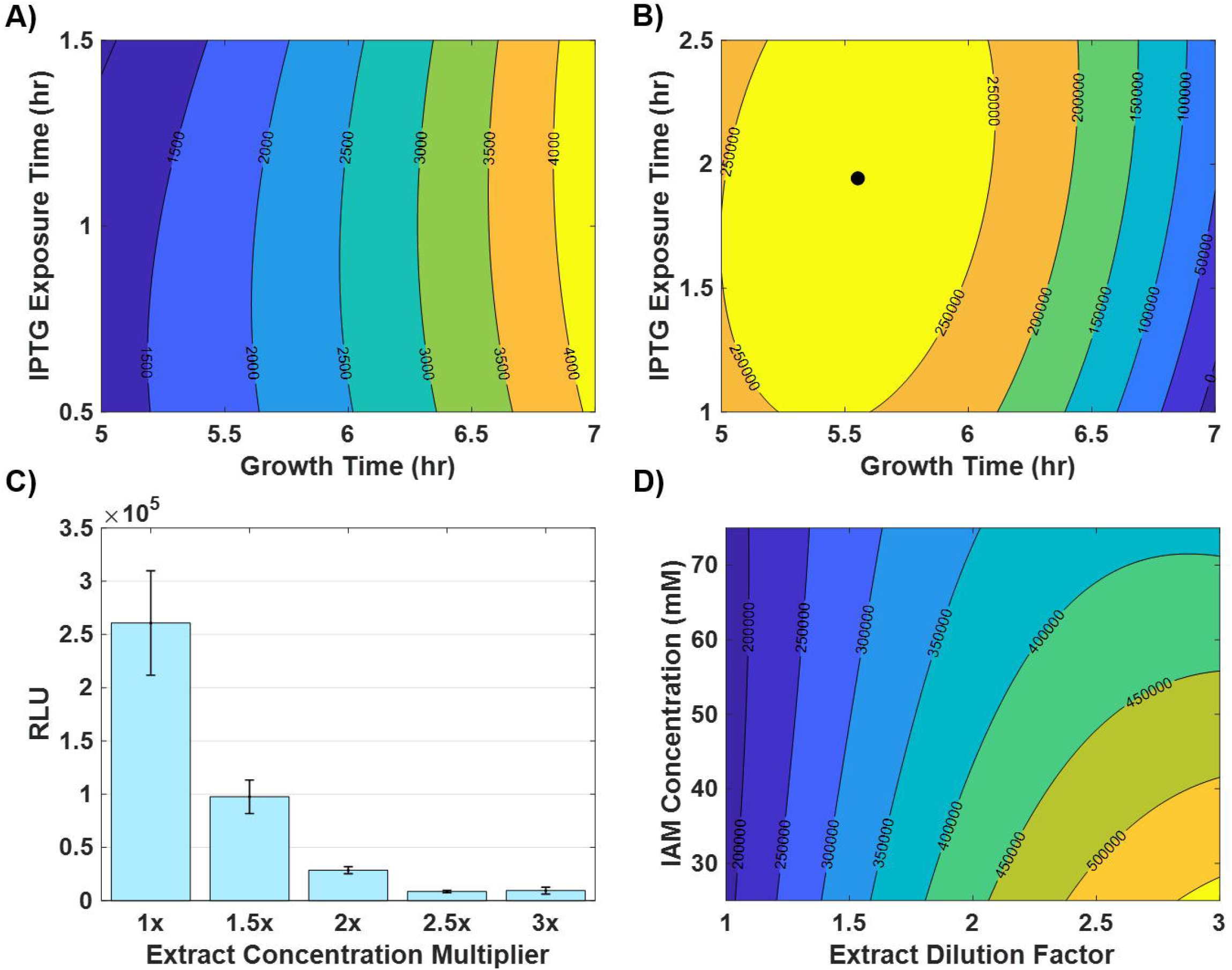
Experiments to optimize expression from T7 Shuffle extract. (A) Response surface fit to determine optimal growth conditions for sfGFP expression; z axis shows fluorescence (B) Response surface fit to determine optimal growth conditions for Gluc expression; z axis shows luminescence.

The response surface for sfGFP shows that longer growth times would improve the expression (Fig 3A). This indicates that insufficient T7 RNAP is being expressed to maximize overall yield; this is also apparent in the level of sfGFP fluorescence observed which is less than we typically observe with BL21 DE3 Star extract when assayed on the same plate. Since sfGFP contains no disulfide bonds, this response surface is not capable of predicting optimum DsbC production for chaperoning disulfide bond formation. This was resolved using our second reporter protein, Gluc. The response surface for Gluc luminescence output has a clear maximum within our experimental bounds (Fig 3B), indicating the optimum balance of chaperone protein and T7 RNAP to produce disulfide proteins. An optimum growth time of 5.6 hr and an optimum induction point at 3.7 hr are found for these smaller growth vessels (200 mL media in 500 mL Tunair flasks), with greater sensitivity in growth time (Supplementary Figure 1A). These optimum induction and harvest time points were then translated to percentages of the total growth curve (fit by a Verhulst-Pearl logistics function) to allow for application to other growth curves occurring in larger vessels or other orbital shakers (50.49% and 13.12% respectively), as we have done before (Dopp and Reuel, 2018). The optimum time points for the 2.5 L shake flasks we use with 1 L culture are then found from these percentages (5.6 and 3.9 hr for harvest and induction respectively). The statistics for both fits are listed in Supplementary Table 2. It should be noted that while Gluc is a convenient reporter protein, optimal extract processing conditions may be protein dependent. This is also evident when using Shuffle for *in vivo* expression conditions where temperature, time of induction, and IPTG concentration must be optimized for each new protein to see best results (Lobstein et al., 2012). However, for the prototyping phase of new proteins we have found sufficient level of expression to quantify protein and measure desired properties using a single general extract; the quest for custom, optimal extracts for each protein is better suited in a later stage such as scale up for production of a few desired proteins.

### Extract and Iodoacetamide Concentration

To improve expression, we explored the impact of extract concentration and adding a common reagent used to inactivate thiol containing reductases, iodoacetamide (IAM) (Kim and Swartz, 2004; Knapp et al., 2007). Previous literature has shown that increasing extract concentration can lead to increased expression rates at the cost of a reduction in total protein produced so we decided to explore the effect of dilution (Didovyk et al., 2017). We assumed that IAM may inhibit the efficacy of the constitutively produced DsbC since IAM is a small molecule that covalently bonds with thiol groups on reductases to reduce their ability to impair disulfide bond formation. Therefore, we reduced the IAM concentration from 50 μM to 30 μM for the experiments shown in Fig 4A. Surprisingly, subsequent dilutions, except 1/5x, outperformed the standard, nondiluted extract. We then tested possible benefits to manipulating the IAM concentration. The previous assumption that IAM would interfere with the thiol groups of endogenous DsbC was observed to be false. Figure 4B shows that of the IAM concentrations screened, 50 μM performed the best. Henceforth, all experiments were conducted with 1/2x, diluted SHuffle extract.

**Figure 4.**
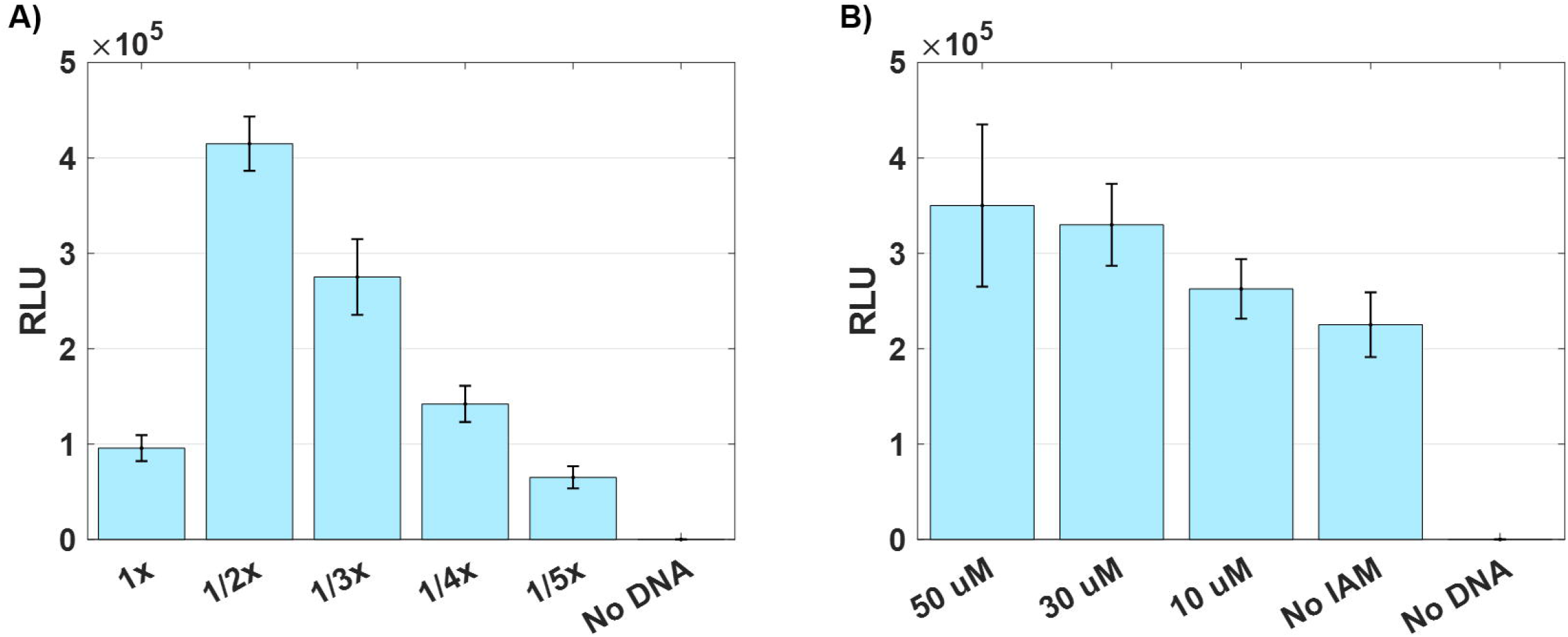
The effects of extract concentration (A) when IAM is held at a constant concentration (30 μM) and the effect of IAM concentration (B) when extract is held at a constant concentration (1/2x) (n=5).

### Production Comparison

To investigate the effect of strain selection on disulfide bond formation, we expressed Gluc using 5 different extracts: SHuffle, KGK10 extract prepared by the Swartz group, KGK10 prepared in our lab, BL21 (DE3) Star, and PURE*frex* 2.1. Figure 5A shows the results of all extracts except PURE*frex* because the signal saturated the detector at gain = 100; by reducing the gain to 80, we can see the superior performance of the commercial kit (Fig 5B). Comparative analysis of the non-commercial extracts is done at gain = 100. The data show that while BL21 (DE3) Star, upon supplementation with DsbC, can support the production of Gluc, the signal is 2.17x lower than that of SHuffle. In contrast, KGK10 extract from the Swartz lab (KGK10 #1) is 1.71x more productive than Shuffle, however, recall that the KGK10 extract requires supplementation of exogenous proteins (T7 RNAP and DsbC) while the Shuffle extract requires neither. Additionally, when we attempt to produce KGK10 using inexpensive shake flasks (Dopp et al., 2019a; Dopp and Reuel, 2018), rather than fermentation in a controlled bioreactor (KGK10 #2), we see very little expression. This suggests that KGK10 requires very strict growth and processing conditions (Zawada and Swartz, 2005) that are inaccessible to labs that do not specialize in cell fermentation (intended ‘users’ of this technology). The SHuffle extract is the best choice for inexpensive, functional *in vitro* production of this reporter protein.

**Figure 5.**
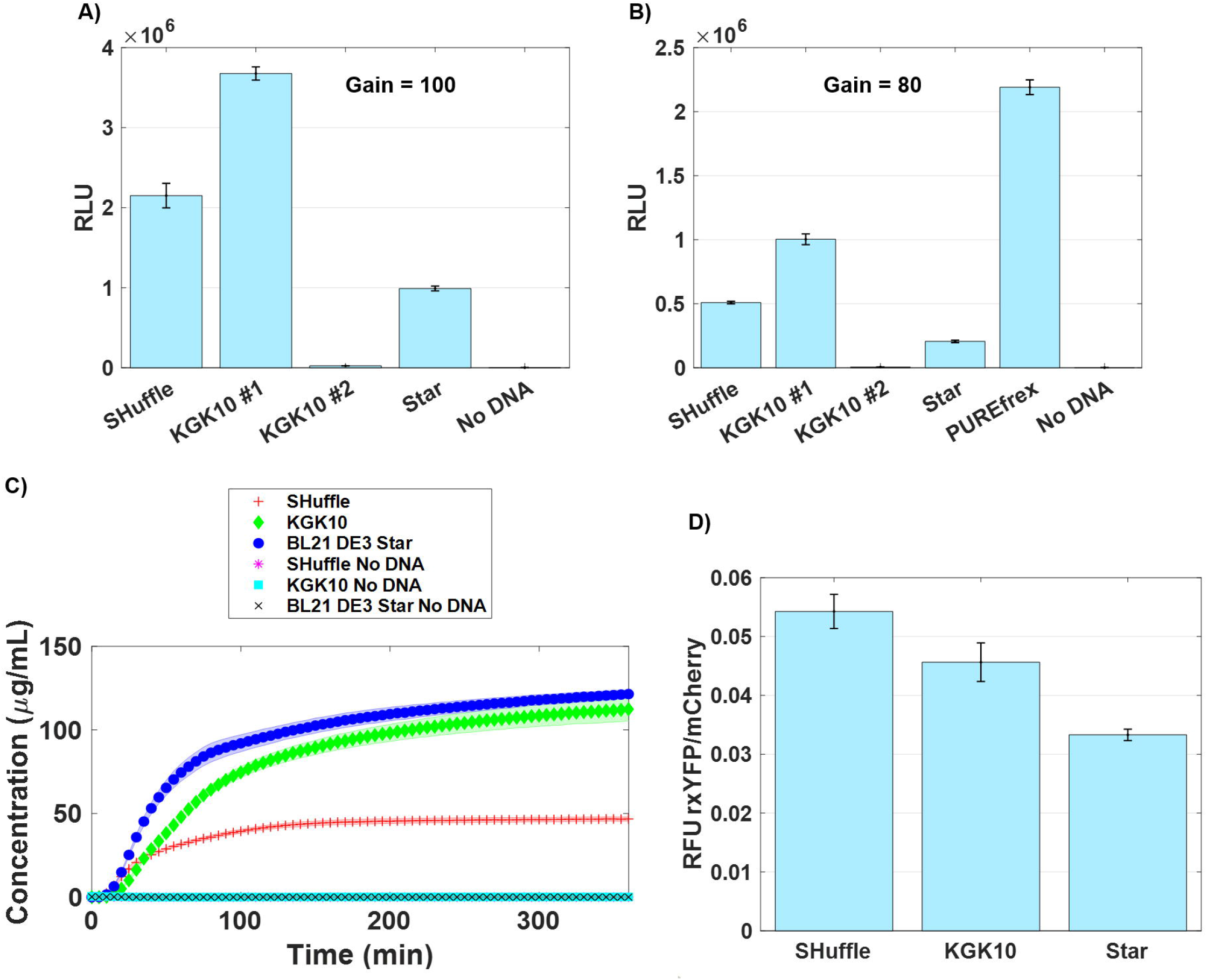
Experiments to compare the yield of T7 SHuffle^®^, KGK10, BL21 DE3 Star, and the PURE*frex* 2.1 kit. KGK10 #1 is carefully fermented KGK10 extract that was provided by the Swartz lab. KGK10 #2 is grown in-house using a shake flask. (A) Gluc signal comparisons at a gain of 100 without PURE*frex* data (n=3). (B) Gluc signal comparisons with an instrument gain of 80 so PURE*frex* data does not saturate the detector. (C) Expression of sfGFP to compare yield of proteins without disulfide bonds over 16 hr (n=3). (D) Is a ratio of oxidation potential/productivity using a YFP-mCherry fusion where a S-S bond has been introduced into a YFP variant (n=3).

Since many proteins of interest do not have disulfide bonds, relative expression levels of proteins without disulfide bonds can also be important information when selecting a strain. To investigate this, we expressed sfGFP using BL21 DE3 Star, KGK10, and T7 SHuffle. We found that BL21 DE3 Star has the most productivity with KGK10 closely following (Fig 5C). This corroborates the finding of our designed experiment which identified the optimal extract for disulfide bond formation to be suboptimal in overall level of protein expression (as measured by sfGFP). These data suggest that the BL21 DE3 star strain is still the better option for proteins with no or few disulfide bonds.

To explore the use of BL21 DE3 Star for proteins with few, proximal disulfide bonds (due to native DsbA present in the extract), we designed a method to compare the ratio between oxidative potential and general productivity. We did this by designing and expressing a new fusion protein made of a yellow fluorescent protein variant (rxYFP) sensitive to the redox environment, a flexible linker, and mCherry to measure overall production. To our knowledge, this is the first fusion protein designed to measure redox/productive potential, especially in cell-free extract. The rxYFP variant has been modified by the addition of two non-native cysteine residues. The fluorescent signal from rxYFP is significantly reduced in the presence of an oxidative environment due to disulfide bond formation between these non-native cysteines (Østergaard et al., 2001). The linker selected was previously used to successfully fuse a nanobody to EGFP and still support proper folding (Li et al., 2012). In order to keep consistent molarity of the DNA template between the reactions, the rxYFP-mCherry fusion template was introduced as linear DNA (see methods). To quantify oxidative potential, the fluorescence of rxYFP was divided by that of mCherry (the lower the ratio, the better the environment for disulfide bond protein expression). Surprisingly, the Shuffle and KGK10 strains have higher ratios than the BL21 DE3, suggesting, again, that the Star strain is suitable to express proteins with low number (in this case 1) of disulfide bonds (Fig 5D).

### Exogenous Supplementation

To explore the impact of added enzymes, exogenous DsbC and T7RNAP were added to the SHuffle reactions. We also investigated the effect of removing these supplements from the fermented KGK10 reactions. Interestingly, the performance of the SHuffle extract decreased when exogenous proteins were added (Fig 6). With the KGK10 extract, we observed that it can still effectively express Gluc without the addition of DsbC, but that T7RNAP is essential for protein production. The tolerance to no chaperone protein may be isolated to this reporter protein as the majority of Gluc’s disulfide bonds occur between consecutive cysteines, thus relying mostly on the native periplasmic isomerase DsbA to form bonds. Previous work with DsbC shows that the bond pattern, or the order in which the cysteines are bonded to one another (consecutive *vs* nonconsecutive), determines how important DsbC is for functional protein production (Berkmen et al., 2005; Ke and Berkmen, 2014; Lobstein et al., 2012). While the KGK10 strain is entirely reliant on exogenous T7RNAP and DsbC for expression, the Shuffle extract’s best expression occurs without the addition of either enzyme. This correlates well with the DoE showing that diluting the extract increases expression in SHuffle. These data again highlight that SHuffle has the advantage of being simpler to use than KGK10. The only requirement for efficient production using SHuffle extract is the optimized induction time of IPTG and the optimized harvest time in a shake flask reactor. KGK10 extract is more complicated to produce because the strain doesn’t produce T7RNAP or DsbC and requires fermentation with a defined media (Knapp et al., 2007).

**Figure 6.**
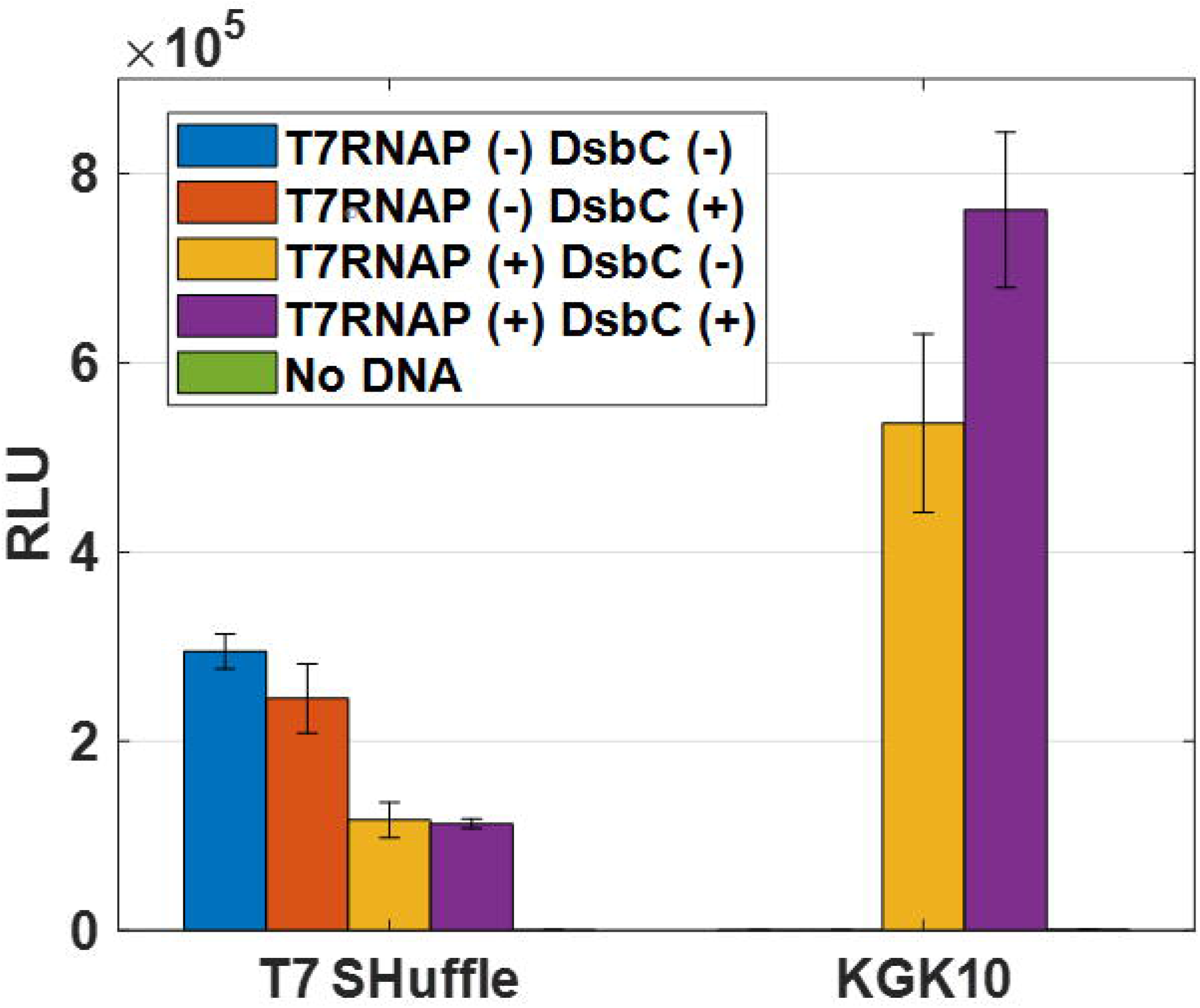
Exploring the effects of including/excluding exogenous T7 RNAP and DsbC to/from cell-free reactions. Gluc activity was measured with luminescence (RLU) (n=3).

### Prototyping Proteins

#### Optimizing Process Conditions

Before prototyping proteins with the Shuffle extract, we determined best process conditions. While typical cell-free reactions are conducted at 37°C for 4 hr, reactions producing proteins with disulfide bonds are commonly conducted at 30°C and conducted overnight (Dopp et al., 2018; Gregorio et al., 2019). The lower temperature has been shown to enhance eukaryotic protein folding efficiency in *E. coli* and longer reaction times can lead to improved titers (Siller et al., 2010). Since we noticed our sfGFP expression flattened off relatively quickly, we decided to test Gluc expression at a shorter time and higher temperature to improve throughput (Fig 7A). The resulting signal is well above what is needed for detection but significantly lower than a 16 hr expression at 30°C. For our work, we have decided to prototype proteins at the slower, but more productive conditions of 16 hr at 30°C. Using these conditions, we then tested the efficacy of different genetic templates at equimolar concentrations (Fig 7B). We’ve previously shown that rolling circle amplification (RCA) can be applied to minimal linear template to avoid time-consuming cloning steps (Dopp et al., 2019b). While the plasmid was much more effective, the signal from the RCA products is still well above noise. We also determined that the SHuffle extract can produce suitable Gluc signal without the 30 min IAM pretreatment in a 4 hr reaction (Supplementary Fig 3). These process conditions were then used for a few prototyping demonstrations, though extract was still pretreated with IAM.

**Figure 7.**
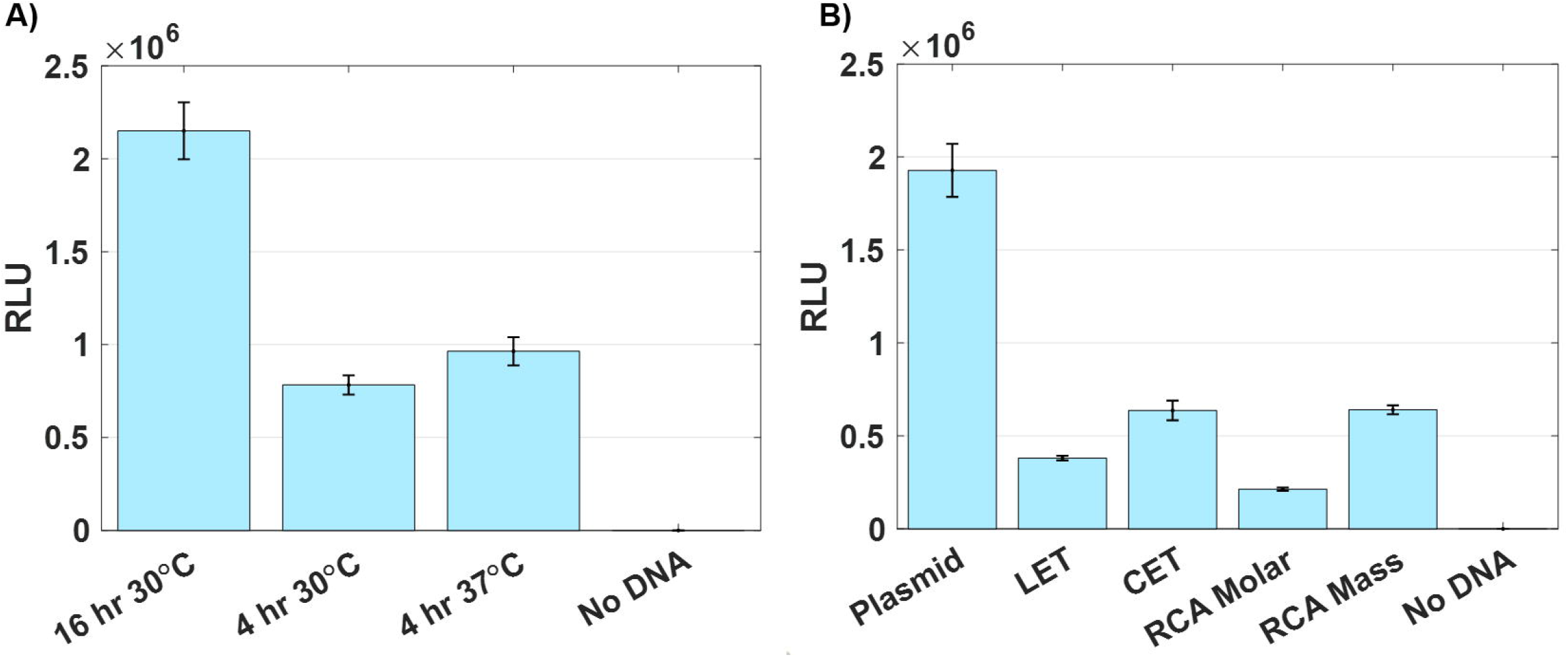
Gluc expressed under conditions that affect screening. (A) The plasmid expressed at different times and temperatures (n=3). (B) Different genetic templates expressed at 30°C for 16 hr (n=3). ‘RCA Molar’ assumes RCA and CET share the same molecular weight so all templates should have an equimolar concentration. ‘RCA Mass’ has the same mass concentration as the plasmid. ‘No DNA’ control shows level of background signal from extract in luminescence assay.

#### Prototyping Luciferases

To demonstrate the prototyping utility of the SHuffle extract, we assembled a panel of 5 luciferase sequences from the Uniprot database; one is the native Gluc sequence while the other four have at least 90% homology with native Gluc but were annotated as ‘uncharacterized’ (never expressed or tested, but found via genome sequencing). A high percentage of homology was chosen to ensure the four, putative luciferases contained multiple disulfide bonds and acted on the substrate coelenterazine. To further demonstrate the power of cell free prototyping, we tested the activity of these putative luciferase enzymes on various coelenterazine analogues to determine if slight differences in amino acid sequence would have noticeable effects on substrate preference (Dikici et al., 2009). As seen in Fig 8, the most effective combination in this panel was the native luciferase from *Gaussia princeps* and the water-soluble analogue of coelenterazine. These results are consistent with current literature (Morse and Tannous, 2012). We can also readily see the substrate compatibility and effect of amino acid sequence changes. Such information could be used in protein engineering studies to improve substrate or protein sequence. All luciferase candidates were expressed from minimal templates that had undergone RCA. The genetic sequences for the luciferase candidates are listed in the Supplementary material (Zhao et al., 2004). Images of the coelenterazine analogues are also provided in the supplement. We were unable to provide a structure for the water soluble (sol) analogue and believe the structure to be proprietary.

**Figure 8.**
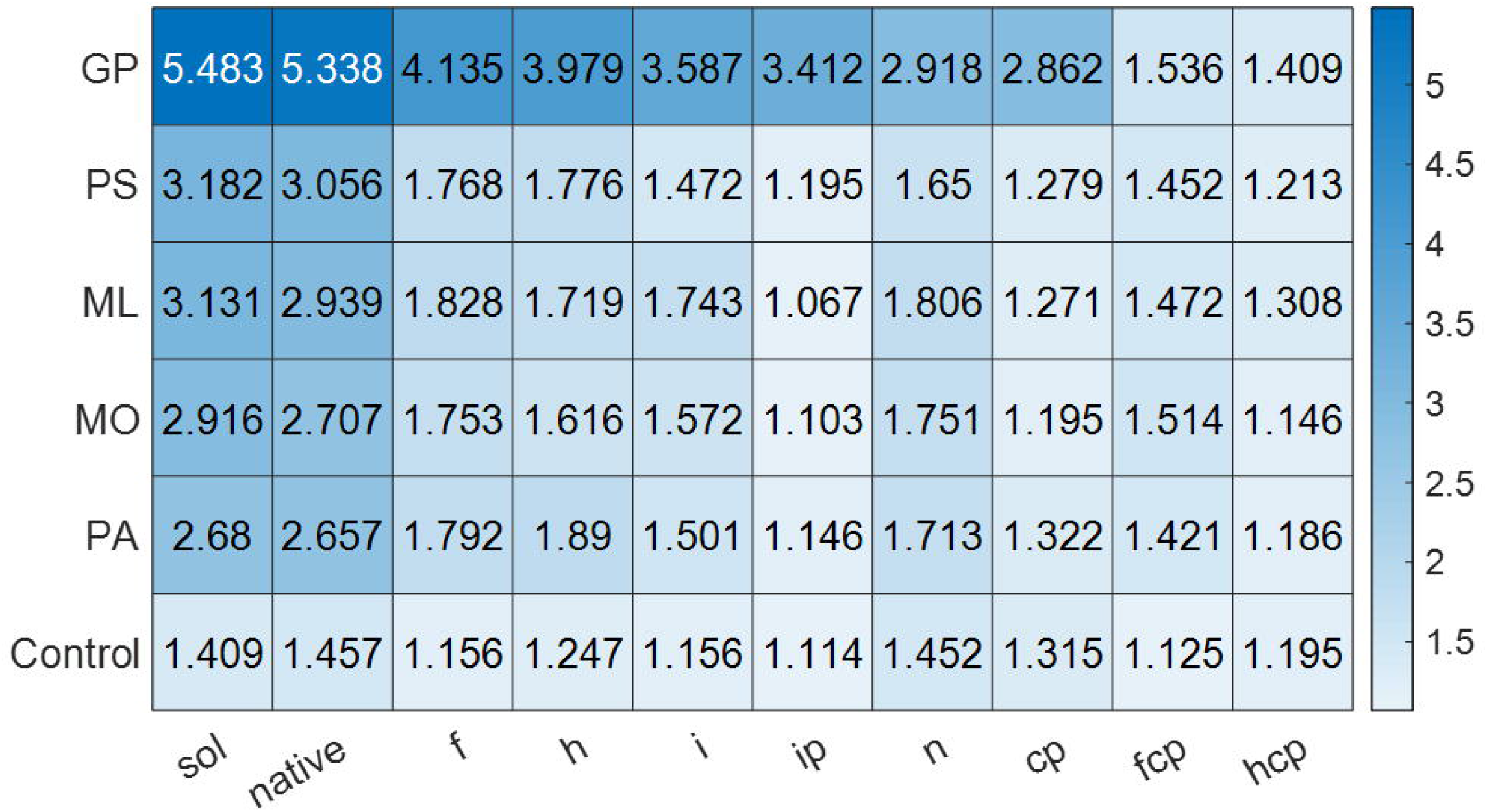
A heat map representing the activity of luciferase from *Gaussia princeps* as well as four other putative luciferases from various organisms against a panel of 10 coelenterazine analogues as the luciferase substrate. The values in the heat map are on a log scale of the recorded luminescence (n=3). GP – *Gaussia princeps* (Uniprot Q9BLZ2), PS - *Pleuromamma scutullata* (Uniprot U3TIM6), ML - *Metridia longa* (Uniprot A0A0B4UFT5), MO - *Metridia okhotensis* (Uniprot H3JS09), PA - *Pleuromamma abdominalis* (Uniprot H3JS12).

#### Prototyping Enzymes

To further investigate the SHuffle strain’s ability to quickly prototype proteins, we chose to express three enzymes (hevamine, endochitinase A (ChitA), and periplasmic AppA (which have 3, 7, and 4 disulfide bonds) also using our previously established minimal, rapid-order genetic template method (Dopp et al., 2019b). Hevamine is a class IIIa chitinase from the rubber tree *Hevea brasiliensis* and exhibits exochitinase activity (Kitaoku et al., 2015). ChitA is a class IV chitinase found primarily in developing maize kernels of *Zea mays* that exhibits endochitinase activity (Volpicella et al., 2017). Both chitinases are responsible for protecting the plants against fungal pathogens (Bokma et al., 2000; Bokma et al., 2002; Hawkins et al., 2015; Volpicella et al., 2017). AppA is a periplasmic protein native to *E. coli* that acts as a phytase (Berkmen et al., 2005). Phytases are an interesting class of enzymes because they break down phytate which is considered an antinutrient and reduces nutrient absorption from seeds (grains, beans, and nuts) in monogastric animals (Bitar and Reinhold, 1972; Zinin et al., 2004). Undigested phytates from monogastric livestock are also a large source of environmental phosphorous pollution in agriculture (Golovan et al., 2001; Zinin et al., 2004). To our knowledge, none of these enzymes have been expressed in an *E. coli*-based cell-free system.

The resulting products can be screened directly in the lysate without purification. Hevamine samples demonstrated 1.3x greater activity than no DNA controls over the length of the entire reaction (Fig 9A). The substrate chose for this reaction was 4-Methylumbelliferyl β-D-N,N′,N″-triacetylchitotrioside which is a fluorogenic used to characterize endochitinases. Hevamine’s kinetics and preferred substrate have been published in previous literature (Bokma et al., 2000). In contrast, ChitA samples performed 2.4x better than the no DNA controls when comparing initial reaction rate (change in fluorescence over time during the first 60 sec, (Fig 9B). The substrate used for ChitA was 4-methylumbelliferyl N-acetyl-β-D-glucosaminide which is fluorogenic substrate used to characterize exochitinases. When screening the expressed AppA, we found the sample with enzyme reaches a kinetic rate of 0.041±.005 abs/min over the span of 5 minutes while the control (no DNA) only reaches 0.031±.005 abs/min (Fig 9C). The substrate used was p-nitrophenyl phosphate which is used to characterize alkaline and acid phosphatase activity.

**Figure 9.**
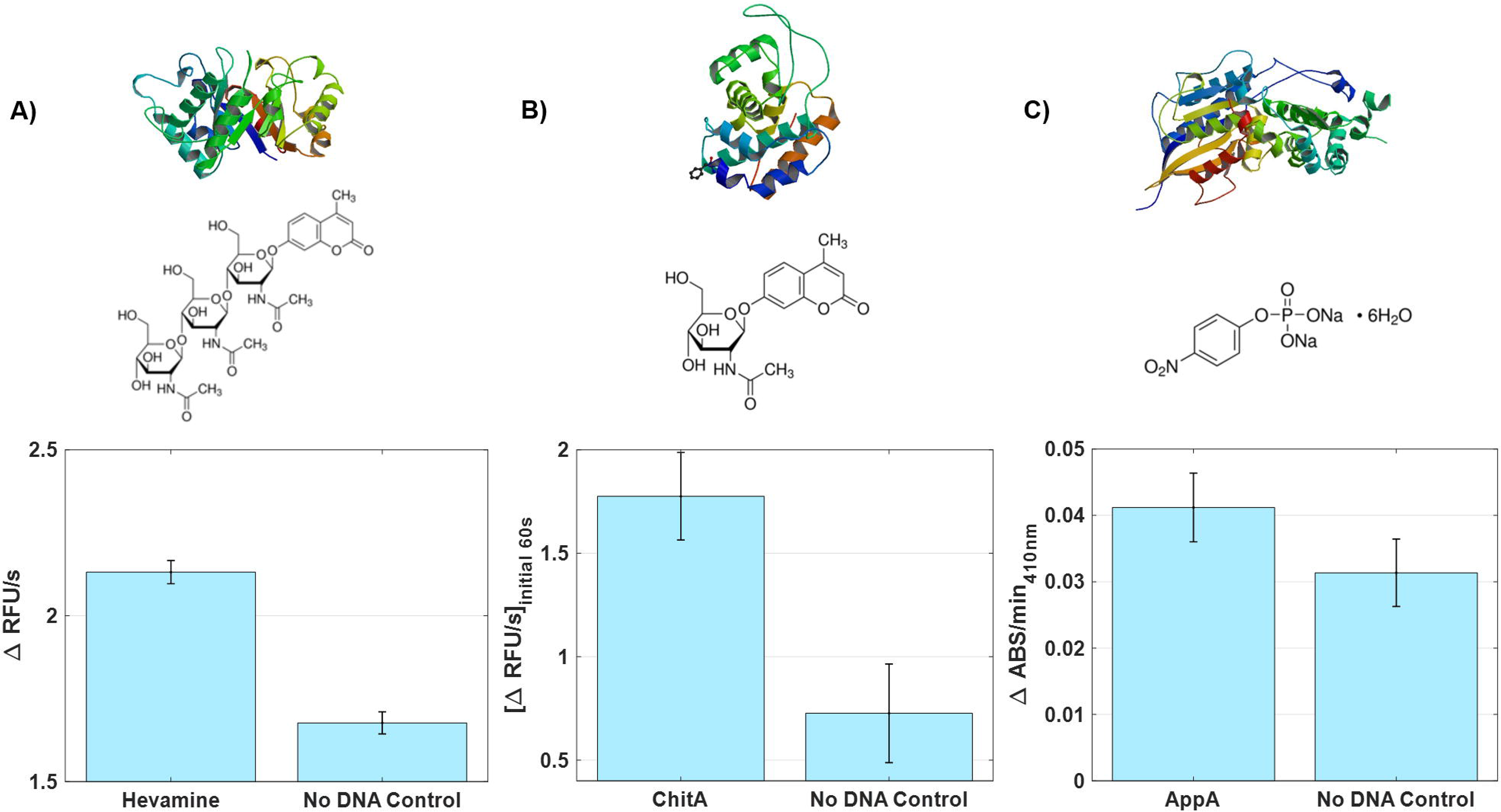
The crystal structure, corresponding substrate, and activity of enzymes used in this work. (A) Total hevamine (RCSB PDB # 2HVM) activity on 4-Methylumbelliferyl β-D-N,N′,N″-triacetylchitotrioside (p-value = 1.10×10^-5^) (n=3). (B) Initial rate (60 sec) of ChitA (RCSB PDB # 4MCK) activity on 4-Methylumbelliferyl N-acetyl-β-D-glucosaminide (p-value = 5.92×10^-4^) (n=4). (C) The activity of AppA (RCSB PDB # 1DKL) on p-nitrophenyl phosphate, disodium salt hexahydrate (p-value = 0.034) (n=4).

## Future Work

We have identified a few strategies to further improve the development and performance of this extract. One source of variability was the use of a reporter protein (Gluc) that requires a secondary assay which has careful, time-based steps to measure level of production (via luminescence). This is much less convenient (and accurate) than measuring the direct signal from a protein rapidly folding (like sfGFP). This could be addressed if a reporter protein is found/designed that can be detected while the reaction is running (such as a fluorescent protein which requires multiple disulfide bonds to produce its fluorescence). Additionally, further genome edits could be made on the SHuffle strain, such as a mutated RNaseE gene (rne131) that makes the Star strain uniquely qualified for expression from LET or RCA templates. Another way to improve expression from linear templates is to add GamS protein to the reaction which can improve expression to near plasmid levels (Ahn et al., 2005; Sitaraman et al., 2004; Sun et al., 2014; Yin et al., 2012). However, this work focuses on the advantage of not adding exogenous proteins. Another strategy of including material, would be the addition of short DNA oligos containing six χ (chi) sites which has been shown to drastically improve yield when using linear DNA (Marshall et al., 2017). This could be included in supplement mix.

## Conclusion

From the experiments performed, we conclude that the commercial cell line T7 SHuffle^®^ can be used to create simple, inexpensive, functional extract for cell free expression of proteins with many disulfide bonds. In processing, this extract is lyophilized to drive off extra water used for the continuous homogenizer which increases the flexibility of extract storage and distribution. This extract is also more economical due to the scalable production method, simple steps and no need for exogenous protein supplementation). This extract can be used for rapid prototyping and design of new proteins from genomic data using minimal genetic templates that are amplified with isothermal rolling circle amplification. In the case of prototyping, where enough protein is needed to measure activity or desired properties, we have demonstrated sufficient level of protein expression. In cases where increased yield is needed, there are some general approaches to further improve yield that we have yet to apply to this strain, such as: increasing the surface area to volume ratio (Gregorio et al., 2019; Hong et al., 2015; Voloshin and Swartz, 2005), implementing machine learning to improve reaction composition (Borkowski et al., 2020), and conducting metabolomic studies to identify and optimize key metabolites (Miguez et al., 2019). However, even with these added improvements, the SHuffle strain may yet express less protein than other strains that are supplemented with exogenous T7RNAP and DsbC, simply due to exhausted machinery. Yet, the simple approach presented in this work can make significant impact at the prototyping phase where a large amount of extract is needed to test many sequence candidates and thus the cost of extract is imperative. Once desired enzymes, therapies, or protein-based materials are discovered, they can be scaled up in more efficient extract or using traditional cell-based methods. Finally, this simple approach which requires less specialized machinery, skill, and time, will be more readily accessible to groups wanting to prototype proteins with disulfide bonds.

## Acknowledgements

The authors acknowledge Prof. Thomas Mansell for the investigative tip to look into T7 SHuffle^®^ capabilities. NFR acknowledges the Black & Veatch Building a World of Difference Faculty Fellow in Engineering for partial funding and Iowa State University Startup Funds. We would also like to thank the Swartz group at Stanford for sharing their knowledge, Gluc plasmid, and KGK10 extract.

