## Supplement for "Simple, Functional, Inexpensive Cell Extract for *in vitro* Prototyping of Proteins with Disulfide Bonds"

### **Supplementary Figures**

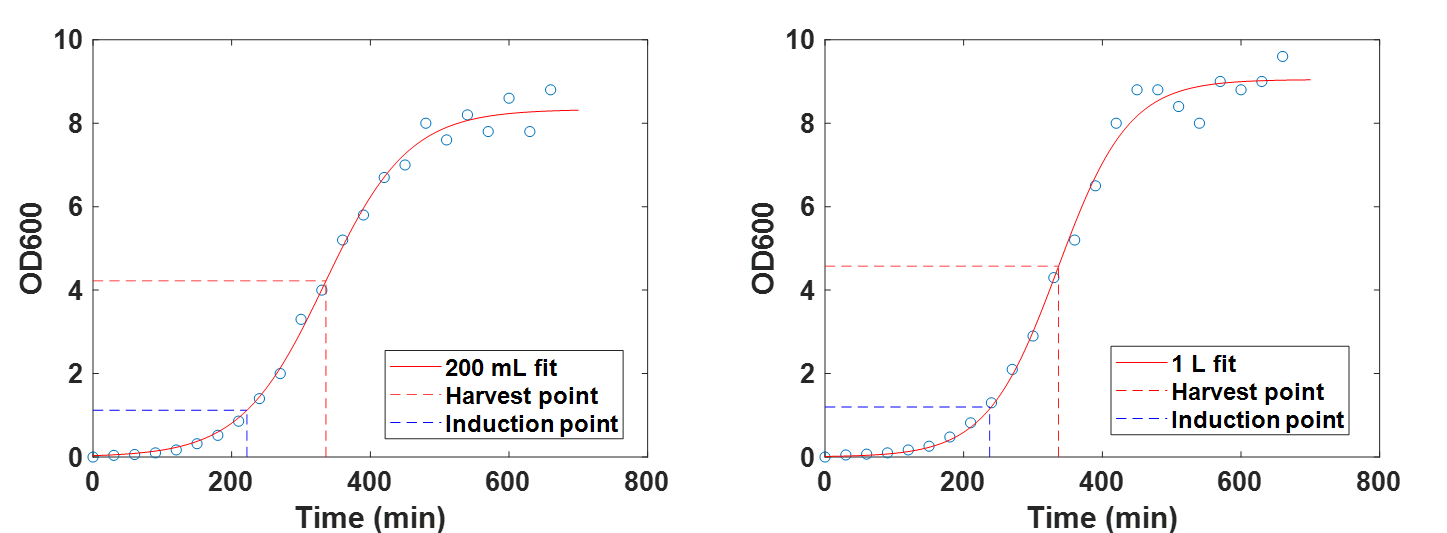

**Supplementary Figure 1** – Growth curves used to correlate data collected from the DoE completed in 500 mL shake flask volumes to 1 L shake flask volumes. (A) The optimum conditions for a 200 mL culture in a 500 mL shake flask inducing with IPTG after 3.7 hrs (222 min) and harvesting after 5.7 hrs (342 min). (B) The optimum time points for a 1 L culture in a 2.5 L shake flask inducing with IPTG after 4.2 hrs (252 min) and harvesting after 5.7 hrs (342 min). These times correspond to 50.49% and 13.12% respectively.

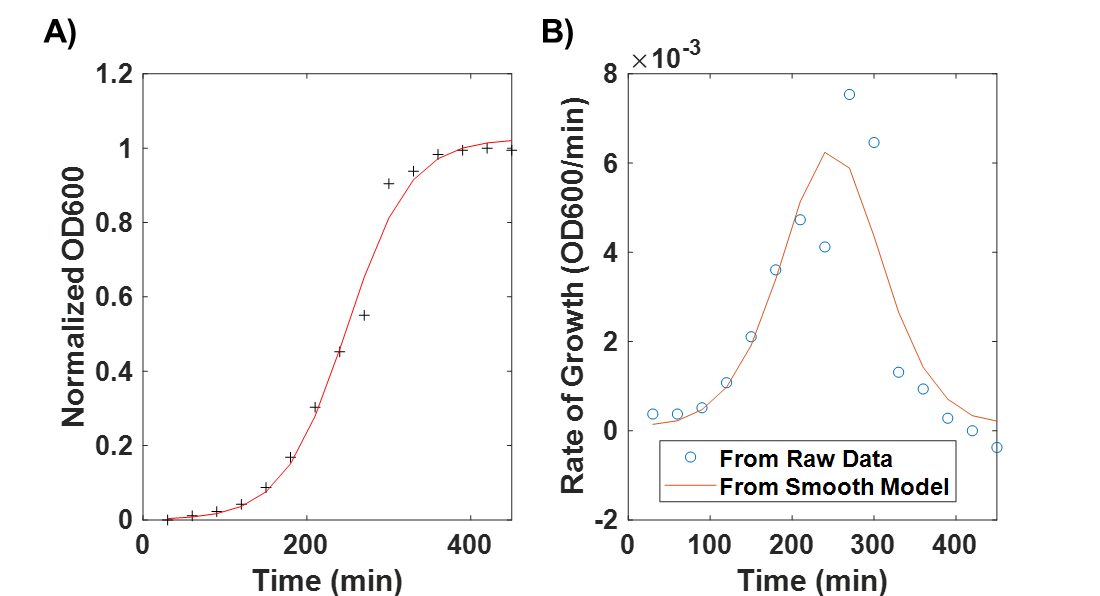

**Supplementary Figure 2** – Plots to determine the optimal harvest point of KGK10. (A) VPE fit of the KGK10 growth curve. (B) First derivative of the growth curve with a model that provides the highest rate of growth at 240 min.

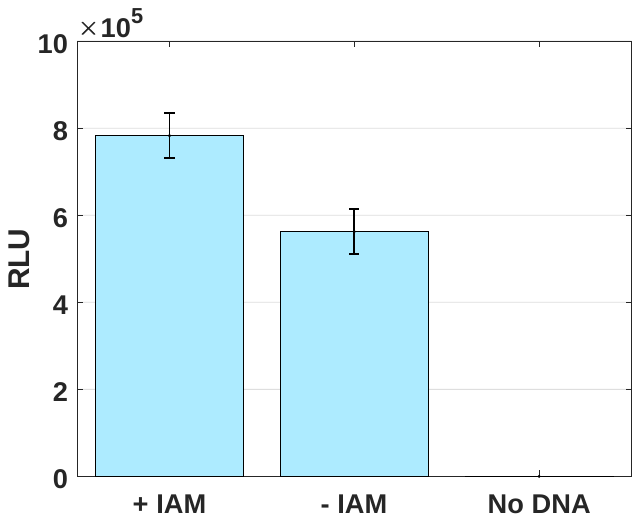

**Supplementary Figure 3** – Plot depicting the performance of SHuffle extract that had been treated with IAM and extract that had not been treated with IAM (n=3).

**Supplementary Table 1** – Time points for the designed experiment to determine the harvest time and the IPTG induction time.

| **Time Points (hr)** | |
| --- | --- |
| **Harvest** | **Induction** |
| 5 | 4 |
| 5 | 3.25 |
| 5 | 2.5 |
| 6 | 5 |
| 6 | 4.25 |
| 6 | 4.25 |
| 6 | 4.25 |
| 6 | 4.25 |
| 6 | 4.25 |
| 6 | 3.5 |
| 7 | 6 |
| 7 | 5.25 |
| 7 | 4.5 |

**Supplementary Table 2** – Statistics obtained from the two DoEs to optimize induction and harvest times using sfGFP and Gluc as reporters, as well as the DoE used to optimize extract and IAM concentrations.

| **Parameter** | **sfGFP** | **Gluc** |
| --- | --- | --- |
| Samples | 13 | 13 |
| RMSE | 631 | 5.7x10^4^ |
| R^2^ | 0.839 | 0.834 |
| Adjusted R^2^ | 0.724 | 0.716 |
| F-Statistic | 7.3 | 7.05 |
| p-value | 0.0106 | 0.0117 |

### Genetic Templates

Below are the genetic templates used CFPS expression. All sequences were optimized for *Escherichia coli* using IDT’s codon optimization tool. The sequences are color coded according to the following key:

T7 Promoter – T7 promoter

RBS – Ribosome binding site

Start – Start codon

Protein Sequence – Gene coding for protein

Stop – Stop codon

T7 Terminator – T7 Terminator

Circularization site – HindIII digest

Primer Sequences – Primer sequences

Linker – Linker connecting rxYFP and mCherry

pET24a-GLuc_6H – 5792 base pairs

TGGCGAATGGGACGCGCCCTGTAGCGGCGCATTAAGCGCGGCGGGTGTGGTGGTTACGCGCAGCGTGACCGCTACACTTGCCAGCGCCCTAGCGCCCGCTCCTTTCGCTTTCTTCCCTTCCTTTCTCGCCACGTTCGCCGGCTTTCCCCGTCAAGCTCTAAATCGGGGGCTCCCTTTAGGGTTCCGATTTAGTGCTTTACGGCACCTCGACCCCAAAAAACTTGATTAGGGTGATGGTTCACGTAGTGGGCCATCGCCCTGATAGACGGTTTTTCGCCCTTTGACGTTGGAGTCCACGTTCTTTAATAGTGGACTCTTGTTCCAAACTGGAACAACACTCAACCCTATCTCGGTCTATTCTTTTGATTTATAAGGGATTTTGCCGATTTCGGCCTATTGGTTAAAAAATGAGCTGATTTAACAAAAATTTAACGCGAATTTTAACAAAATATTAACGTTTACAATTTCAGGTGGCACTTTTCGGGGAAATGTGCGCGGAACCCCTATTTGTTTATTTTTCTAAATACATTCAAATATGTATCCGCTCATGAATTAATTCTTAGAAAAACTCATCGAGCATCAAATGAAACTGCAATTTATTCATATCAGGATTATCAATACCATATTTTTGAAAAAGCCGTTTCTGTAATGAAGGAGAAAACTCACCGAGGCAGTTCCATAGGATGGCAAGATCCTGGTATCGGTCTGCGATTCCGACTCGTCCAACATCAATACAACCTATTAATTTCCCCTCGTCAAAAATAAGGTTATCAAGTGAGAAATCACCATGAGTGACGACTGAATCCGGTGAGAATGGCAAAAGTTTATGCATTTCTTTCCAGACTTGTTCAACAGGCCAGCCATTACGCTCGTCATCAAAATCACTCGCATCAACCAAACCGTTATTCATTCGTGATTGCGCCTGAGCGAGACGAAATACGCGATCGCTGTTAAAAGGACAATTACAAACAGGAATCGAATGCAACCGGCGCAGGAACACTGCCAGCGCATCAACAATATTTTCACCTGAATCAGGATATTCTTCTAATACCTGGAATGCTGTTTTCCCGGGGATCGCAGTGGTGAGTAACCATGCATCATCAGGAGTACGGATAAAATGCTTGATGGTCGGAAGAGGCATAAATTCCGTCAGCCAGTTTAGTCTGACCATCTCATCTGTAACATCATTGGCAACGCTACCTTTGCCATGTTTCAGAAACAACTCTGGCGCATCGGGCTTCCCATACAATCGATAGATTGTCGCACCTGATTGCCCGACATTATCGCGAGCCCATTTATACCCATATAAATCAGCATCCATGTTGGAATTTAATCGCGGCCTAGAGCAAGACGTTTCCCGTTGAATATGGCTCATAACACCCCTTGTATTACTGTTTATGTAAGCAGACAGTTTTATTGTTCATGACCAAAATCCCTTAACGTGAGTTTTCGTTCCACTGAGCGTCAGACCCCGTAGAAAAGATCAAAGGATCTTCTTGAGATCCTTTTTTTCTGCGCGTAATCTGCTGCTTGCAAACAAAAAAACCACCGCTACCAGCGGTGGTTTGTTTGCCGGATCAAGAGCTACCAACTCTTTTTCCGAAGGTAACTGGCTTCAGCAGAGCGCAGATACCAAATACTGTCCTTCTAGTGTAGCCGTAGTTAGGCCACCACTTCAAGAACTCTGTAGCACCGCCTACATACCTCGCTCTGCTAATCCTGTTACCAGTGGCTGCTGCCAGTGGCGATAAGTCGTGTCTTACCGGGTTGGACTCAAGACGATAGTTACCGGATAAGGCGCAGCGGTCGGGCTGAACGGGGGGTTCGTGCACACAGCCCAGCTTGGAGCGAACGACCTACACCGAACTGAGATACCTACAGCGTGAGCTATGAGAAAGCGCCACGCTTCCCGAAGGGAGAAAGGCGGACAGGTATCCGGTAAGCGGCAGGGTCGGAACAGGAGAGCGCACGAGGGAGCTTCCAGGGGGAAACGCCTGGTATCTTTATAGTCCTGTCGGGTTTCGCCACCTCTGACTTGAGCGTCGATTTTTGTGATGCTCGTCAGGGGGGCGGAGCCTATGGAAAAACGCCAGCAACGCGGCCTTTTTACGGTTCCTGGCCTTTTGCTGGCCTTTTGCTCACATGTTCTTTCCTGCGTTATCCCCTGATTCTGTGGATAACCGTATTACCGCCTTTGAGTGAGCTGATACCGCTCGCCGCAGCCGAACGACCGAGCGCAGCGAGTCAGTGAGCGAGGAAGCGGAAGAGCGCCTGATGCGGTATTTTCTCCTTACGCATCTGTGCGGTATTTCACACCGCATATATGGTGCACTCTCAGTACAATCTGCTCTGATGCCGCATAGTTAAGCCAGTATACACTCCGCTATCGCTACGTGACTGGGTCATGGCTGCGCCCCGACACCCGCCAACACCCGCTGACGCGCCCTGACGGGCTTGTCTGCTCCCGGCATCCGCTTACAGACAAGCTGTGACCGTCTCCGGGAGCTGCATGTGTCAGAGGTTTTCACCGTCATCACCGAAACGCGCGAGGCAGCTGCGGTAAAGCTCATCAGCGTGGTCGTGAAGCGATTCACAGATGTCTGCCTGTTCATCCGCGTCCAGCTCGTTGAGTTTCTCCAGAAGCGTTAATGTCTGGCTTCTGATAAAGCGGGCCATGTTAAGGGCGGTTTTTTCCTGTTTGGTCACTGATGCCTCCGTGTAAGGGGGATTTCTGTTCATGGGGGTAATGATACCGATGAAACGAGAGAGGATGCTCACGATACGGGTTACTGATGATGAACATGCCCGGTTACTGGAACGTTGTGAGGGTAAACAACTGGCGGTATGGATGCGGCGGGACCAGAGAAAAATCACTCAGGGTCAATGCCAGCGCTTCGTTAATACAGATGTAGGTGTTCCACAGGGTAGCCAGCAGCATCCTGCGATGCAGATCCGGAACATAATGGTGCAGGGCGCTGACTTCCGCGTTTCCAGACTTTACGAAACACGGAAACCGAAGACCATTCATGTTGTTGCTCAGGTCGCAGACGTTTTGCAGCAGCAGTCGCTTCACGTTCGCTCGCGTATCGGTGATTCATTCTGCTAACCAGTAAGGCAACCCCGCCAGCCTAGCCGGGTCCTCAACGACAGGAGCACGATCATGCGCACCCGTGGGGCCGCCATGCCGGCGATAATGGCCTGCTTCTCGCCGAAACGTTTGGTGGCGGGACCAGTGACGAAGGCTTGAGCGAGGGCGTGCAAGATTCCGAATACCGCAAGCGACAGGCCGATCATCGTCGCGCTCCAGCGAAAGCGGTCCTCGCCGAAAATGACCCAGAGCGCTGCCGGCACCTGTCCTACGAGTTGCATGATAAAGAAGACAGTCATAAGTGCGGCGACGATAGTCATGCCCCGCGCCCACCGGAAGGAGCTGACTGGGTTGAAGGCTCTCAAGGGCATCGGTCGAGATCCCGGTGCCTAATGAGTGAGCTAACTTACATTAATTGCGTTGCGCTCACTGCCCGCTTTCCAGTCGGGAAACCTGTCGTGCCAGCTGCATTAATGAATCGGCCAACGCGCGGGGAGAGGCGGTTTGCGTATTGGGCGCCAGGGTGGTTTTTCTTTTCACCAGTGAGACGGGCAACAGCTGATTGCCCTTCACCGCCTGGCCCTGAGAGAGTTGCAGCAAGCGGTCCACGCTGGTTTGCCCCAGCAGGCGAAAATCCTGTTTGATGGTGGTTAACGGCGGGATATAACATGAGCTGTCTTCGGTATCGTCGTATCCCACTACCGAGATATCCGCACCAACGCGCAGCCCGGACTCGGTAATGGCGCGCATTGCGCCCAGCGCCATCTGATCGTTGGCAACCAGCATCGCAGTGGGAACGATGCCCTCATTCAGCATTTGCATGGTTTGTTGAAAACCGGACATGGCACTCCAGTCGCCTTCCCGTTCCGCTATCGGCTGAATTTGATTGCGAGTGAGATATTTATGCCAGCCAGCCAGACGCAGACGCGCCGAGACAGAACTTAATGGGCCCGCTAACAGCGCGATTTGCTGGTGACCCAATGCGACCAGATGCTCCACGCCCAGTCGCGTACCGTCTTCATGGGAGAAAATAATACTGTTGATGGGTGTCTGGTCAGAGACATCAAGAAATAACGCCGGAACATTAGTGCAGGCAGCTTCCACAGCAATGGCATCCTGGTCATCCAGCGGATAGTTAATGATCAGCCCACTGACGCGTTGCGCGAGAAGATTGTGCACCGCCGCTTTACAGGCTTCGACGCCGCTTCGTTCTACCATCGACACCACCACGCTGGCACCCAGTTGATCGGCGCGAGATTTAATCGCCGCGACAATTTGCGACGGCGCGTGCAGGGCCAGACTGGAGGTGGCAACGCCAATCAGCAACGACTGTTTGCCCGCCAGTTGTTGTGCCACGCGGTTGGGAATGTAATTCAGCTCCGCCATCGCCGCTTCCACTTTTTCCCGCGTTTTCGCAGAAACGTGGCTGGCCTGGTTCACCACGCGGGAAACGGTCTGATAAGAGACACCGGCATACTCTGCGACATCGTATAACGTTACTGGTTTCACATTCACCACCCTGAATTGACTCTCTTCCGGGCGCTATCATGCCATACCGCGAAAGGTTTTGCGCCATTCGATGGTGTCCGGGATCTCGACGCTCTCCCTTATGCGACTCCTGCATTAGGAAGCAGCCCAGTAGTAGGTTGAGGCCGTTGAGCACCGCCGCCGCAAGGAATGGTGCATGCAAGGAGATGGCGCCCAACAGTCCCCCGGCCACGGGGCCTGCCACCATACCCACGCCGAAACAAGCGCTCATGAGCCCGAAGTGGCGAGCCCGATCTTCCCCATCGGTGATGTCGGCGATATAGGCGCCAGCAACCGCACCTGTGGCGCCGGTGATGCCGGCCACGATGCGTCCGGCGTAGAGGATCGAGATCTCGATCCCGCGAAATTAATACGACTCACTATAGGGGAATTGTGAGCGGATAACAATTCCCCTCTAGAAATAATTTTGTTTAACTTTAAGAAGGAGATATACATATGGCGAAACCAACAGAAAATAACGAAGACTTCAACATCGTGGCCGTGGCCAGCAACTTCGCGACCACGGATCTCGATGCTGACCGCGGGAAGTTGCCCGGCAAGAAGCTGCCGCTGGAGGTGCTCAAAGAGCTGGAAGCCAATGCCCGGAAAGCTGGCTGCACCAGGGGCTGTCTGATCTGCCTGTCCCACATCAAGTGCACGCCCAAGCTGAAGAAGTTCATCCCAGGACGCTGCCACACCTACGAAGGCGACAAAGAGTCCGCACAGGGCGGCATAGGCGAGGCGATCGTCGACATTCCTGAGATTCCTGGGTTCAAGGACTTGGAGCCCCTGGAGCAGTTCATCGCACAGGTCGATCTGTGTGTGGACTGCACAACTGGCTGCCTCAAAGGGCTTGCCAACGTGCAGTGTTCTGACCTGCTCAAGAAGTGGCTGCCGCAACGCTGTGCGACCTTTGCCAGCAAGATCCAGGGCCAGGTGGACAAGATCAAGGGGGCCGGTGGTGACATGCATCACCATCACCATCACTAATAAGTCGACAAGCTTGCGGCCGCACTCGAGCACCACCACCACCACCACTGAGATCCGGCTGCTAACAAAGCCCGAAAGGAAGCTGAGTTGGCTGCTGCCACCGCTGAGCAATAACTAGCATAACCCCTTGGGGCCTCTAAACGGGTCTTGAGGGGTTTTTTGCTGAAAGGAGGAACTATATCCGGAT

*Gaussia Princeps* Luciferase – 792 base pairs

GTAAAACGACGGCCAGTAGCGCTATTAAAGCTTcgaaatTAATACGACTCACTATAGGGAGACCACAACGGTTTCCCTCTAGAAATAATTTTGTTTAACTTTAAGAAGGAGATATACATATGAAGCCCACGGAGAATAATGAAGACTTTAACATAGTGGCAGTCGCATCGAACTTTGCTACAACTGATCTTGATGCTGATCGGGGTAAGTTACCAGGGAAAAAACTTCCGTTAGAAGTATTAAAGGAAATGGAGGCCAACGCGCGCAAGGCGGGATGTACCAGAGGCTGTCTGATATGCTTGTCGCACATAAAGTGTACGCCCAAAATGAAAAAGTTCATCCCAGGACGCTGTCATACGTACGAAGGCGATAAAGAGTCAGCTCAAGGGGGCATAGGAGAAGCTATTGTAGACATCCCGGAAATACCAGGTTTCAAAGATTTAGAACCGATGGAGCAGTTTATAGCCCAAGTGGATTTGTGCGTTGATTGTACGACAGGGTGTCTTAAAGGACTTGCAAACGTGCAGTGCAGCGACTTATTGAAGAAATGGTTGCCTCAACGGTGCGCCACTTTTGCCTCAAAAATCCAAGGCCAGGTTGATAAGATAAAGGGAGCCGGGGGTGACTAATAAGTCGACCGGCTGCTAACAAAGCCCGAAAGGAAGCTGAGTTGGCTGCTGCCACCGCTGAGCAATAACTAGCATAACCCCTTGGGGCCTCTAAACGGGTCTTGAGGGGTTTTTTGCTGAAAGCGAGACTAAGCTTTAAACTTCGGGTCATAGCTGTTTCCTG

*Metridia longa* (ML) Luciferase – 792 base pairs

GTAAAACGACGGCCAGTAGCGCTATTAAAGCTTcgaaatTAATACGACTCACTATAGGGAGACCACAACGGTTTCCCTCTAGAAATAATTTTGTTTAACTTTAAGAAGGAGATATACATATGGACATTAAATTCATTTTCGCATTGGTATGCATAGCCTTGGTTCAAGCAAACCCTACCGTCAATAATGACGTGAATCGTGGGAAGATGCCGGGTAAGAAATTACCCCTTGAGGTATTGATCGAAATGGAGGCGAATGCCTTCAAGGCAGGATGTACCCGTGGGTGCCTTATTTGCTTGTCGAAGATTAAATGCACGGCCAAAATGAAACAATATATACCTGGGCGCTGCCACGATTATGGGGGCGATAAGAAAACGGGGCAAGCAGGCATTGTGGGCGCAATAGTCGATATTCCCGAGATCAGCGGCTTCAAAGAAATGGAGCCTATGGAGCAATTTATTGCTCAAGTCGATCTGTGCGCTGACTGTACCACGGGCTGCTTGAAAGGTTTAGCTAACGTGAAGTGCTCGGAGTTATTGAAAAAATGGCTTCCTGATCGCTGTGCATCTTTCGCGGATAAGATACAAAAAGAAGCGCACAATATCAAAGGTCTGGCAGGGGACCGCTAATAAGTCGACCGGCTGCTAACAAAGCCCGAAAGGAAGCTGAGTTGGCTGCTGCCACCGCTGAGCAATAACTAGCATAACCCCTTGGGGCCTCTAAACGGGTCTTGAGGGGTTTTTTGCTGAAAGCGAGACTAAGCTTTAAACTTCGGGTCATAGCTGTTTCCTG

*Pleuromamma abdominalis* (PA)Luciferase – 843 base pairs

GTAAAACGACGGCCAGTAGCGCTATTAAAGCTTcgaaatTAATACGACTCACTATAGGGAGACCACAACGGTTTCCCTCTAGAAATAATTTTGTTTAACTTTAAGAAGGAGATATACATATGGCCCTGAAATTCTTGGTCGCAGTAATCTGTTTAGCGGCGGTGCAAGCCAAGTCGATCGACAGTTACGAGAACATTGATATTGTAGCCGTAGCAGGGAACTTTGCGGCGGTCGACCAGGACGCGAACCGGGGTGGAAACCTGCCCGGTAAAAAGATGCCCATTGAAGTCCTTAAAGAGATGGAAGCCAACGCGAAACGGGCGGGCTGCGTTAGAGGTTGCTTGATTTGCTTATCACATATTAAATGTACGGCTAAAATGAAGAAGTTTATTCCGGGCCGTTGTCATTCATATCACGGGGATGCCGACACCAAGCAAGGAGCCCTGGAGGAAGTTGTTGATATGCCTGAAATCCCAGGCTTTGTCGACATGGAGCCGATGGAACAGTTTATAGCGCAGGTCGATAAGTGCGAAGACTGTACTACAGGGTGTTTGAAGGGGTTAGCGAATGTGCACTGCAGTGATCTTCTTAAAAAATGGTTGCCACAGCGGTGCTCCCAATTTGCTGACAAGATACAATCAGAGGTTGACACTATCAAGGGTTTAGCCGGTGACAGATAATAAGTCGACCGGCTGCTAACAAAGCCCGAAAGGAAGCTGAGTTGGCTGCTGCCACCGCTGAGCAATAACTAGCATAACCCCTTGGGGCCTCTAAACGGGTCTTGAGGGGTTTTTTGCTGAAAGCGAGACTAAGCTTTAAACTTCGGGTCATAGCTGTTTCCTG

*Metridia okhotensis* (MO) Luciferase – 936 base pairs

GTAAAACGACGGCCAGTAGCGCTATTAAAGCTTcgaaatTAATACGACTCACTATAGGGAGACCACAACGGTTTCCCTCTAGAAATAATTTTGTTTAACTTTAAGAAGGAGATATACATATGCCGCGCGGCAATATGGACATTAAAGTCCTTTTTGCTCTGACTTGTTTCGCGCTTGTGCAGTCGAACCCGACCGAGACACAGGATGGAGTCGACATTTTAGGAGTAGAGGGGAAATTCGGAACAGAAACCAACCTGGAAACTGACTTGTTTACCATCTGGGAGATTAACGGCATTATAAAATCGGATAGAGACACGAACCGCGCCAACACTGACGCAGACAGAGGTAAGATGCCTGGTAAAAAGTTACCTTTAGCGGTTTTAATAGAAATGGAGGCGAACGCCTTCAAAGCAGGCTGCACCCGTGGTTGCTTAATCTGCCTTAGCAAGATTAAATGCACCGCCAAAATGAAGGAATATATACCTGGAAGATGTCACGATTATGGAGGTGACAAGAAGACAGGGCAGGCGGGAATAGTTGGAGCCATCGTGGATATTCCAGAAATCTCGGGTTTTAAGGAACTGGGGCCAATGGAGCAGTTTATCGCTCAAGTAGATCTGTGCGCCGATTGCACGACGGGCTGCTTAAAAGGTCTTGCTAATGTAAAATGCTCGGCCTTGTTGAAAAAATGGTTACCGGATAGATGTGCGAGTTTCGCTGATAAAATACAGCGCGAAGTGCACAATATCAAGGGCTTGGCGGGTGATCGCTAATAAGTCGACCGGCTGCTAACAAAGCCCGAAAGGAAGCTGAGTTGGCTGCTGCCACCGCTGAGCAATAACTAGCATAACCCCTTGGGGCCTCTAAACGGGTCTTGAGGGGTTTTTTGCTGAAAGCGAGACTAAGCTTTAAACTTCGGGTCATAGCTGTTTCCTG

*Pleuromamma scutullata* (PS) Luciferase – 945 base pairs

GTAAAACGACGGCCAGTAGCGCTATTAAAGCTTcgaaatTAATACGACTCACTATAGGGAGACCACAACGGTTTCCCTCTAGAAATAATTTTGTTTAACTTTAAGAAGGAGATATACATATGTCTATTCAGTTCTTATATGCGTTGGTCTGCTTAGCCGCAGCCGGCTGTCAGTCGCAGAAGTTGTTGCCCTCAGAAGATCCTGAACAGTATAACATTGCGGACCTTGATGACTTGGTTGCCAAGTTGAGTATAACTGACGACGAGATGGAAACCTATACCATCTGGGAAGAACTGCTTATTATTAGTCAGGATTTTGCAAACAACTTGAATGTCGTGGACGGGGACAGAGATCGTAAGTTGCCAGGGAAAAAACTGCCTCTGGAAGTGCTGAAGATCATGGAAGCCAATGCGCGCCGGGCGGGGTGTACTCGGGGCTGCTTAATTTGCTTATCGAAAATCAAGTGTACCGCAAAAATGAAGAAGTTTATTCCAGGTCGTTGCCACACATACGAAGGTGATAAGTCCATTGGTCAAGGTGGCATTGGGGCGGCTATTATCGATATACCAGAAATTCCTGGTTTCAAGGAATTGGAACCTATGGAGCAATTCATTGCACAAGTAGATTTGTGTGCAGATTGTACGACAAGATGCTTAAAAGGCTTGGCAAATGTGCGCTGCAACGATCTGTTAAAAAAATGGTTGCCTGACAGATGTGCGGGCTTCGCGAACAAAATACAATCTGAAGTCCACAACATCAAGGGTCTGGCGGGCGACCGGTAATAAGTCGACCGGCTGCTAACAAAGCCCGAAAGGAAGCTGAGTTGGCTGCTGCCACCGCTGAGCAATAACTAGCATAACCCCTTGGGGCCTCTAAACGGGTCTTGAGGGGTTTTTTGCTGAAAGCGAGACTAAGCTTTAAACTTCGGGTCATAGCTGTTTCCTG

6x His-Hevamine – 1161 base pairs

GTAAAACGACGGCCAGTAGCGCTATTAAAGCTTCGAAATTAATACGACTCACTATAGGGAGACCACAACGGTTTCCCTCTAGAAATAATTTTGTTTAACTTTAAGAAGGAGATATACATATGCACCACCACCATCATCACGGGGGAATCGCAATCTATTGGGGACAAAACGGTAACGAAGGCACGCTTACCCAGACCTGTAGCACTCGGAAATATAGCTACGTAAACATAGCGTTTCTGAACAAATTTGGGAACGGACAGACGCCTCAGATCAACCTGGCGGGACATTGTAATCCAGCCGCAGGCGGTTGTACAATAGTGTCCAATGGTATACGTTCGTGCCAAATTCAAGGGATCAAGGTCATGTTGAGTTTGGGAGGTGGAATAGGATCATATACCCTGGCAAGCCAAGCAGACGCTAAAAACGTGGCCGATTATTTGTGGAACAATTTCCTGGGTGGTAAGAGTAGTTCCCGTCCATTGGGGGACGCTGTCTTAGATGGGATCGACTTTGATATCGAGCATGGGTCTACTCTGTACTGGGACGACCTTGCTCGTTATTTGTCAGCCTACTCTAAACAAGGGAAAAAGGTGTACTTGACAGCAGCTCCGCAGTGTCCCTTTCCCGACCGCTATTTAGGCACGGCCTTAAATACCGGCCTGTTTGATTATGTTTGGGTGCAATTCTACAACAACCCCCCTTGTCAATACTCAAGTGGAAATATAAACAACATTATCAACAGTTGGAACCGTTGGACAACCAGTATAAACGCTGGTAAGATATTTCTTGGGTTGCCAGCAGCACCGGAAGCTGCTGGGTCTGGTTATGTGCCACCGGACGTGCTTATATCTAGAATTCTTCCAGAGATCAAAAAGTCTCCAAAGTATGGCGGTGTAATGTTATGGTCAAAGTTCTATGACGATAAAAATGGCTACTCCAGTTCAATTCTTGATTCGGTCTTGTTTTTACACAGCGAGGAGTGTATGACCGTATTATAATAAGTCGACCGGCTGCTAACAAAGCCCGAAAGGAAGCTGAGTTGGCTGCTGCCACCGCTGAGCAATAACTAGCATAACCCCTTGGGGCCTCTAAACGGGTCTTGAGGGGTTTTTTGCTGAAAGCGAGACTAAGCTTTAAACTTCGGGTCATAGCTGTTTCCTG

6x His-Endochitinase A – 1071 base pairs

GTAAAACGACGGCCAGTAGCGCTATTAAAGCTTCGAAATTAATACGACTCACTATAGGGAGACCACAACGGTTTCCCTCTAGAAATAATTTTGTTTAACTTTAAGAAGGAGATATACATATGCACCATCACCATCATCATCAGAATTGCGGCTGCCAACCAAATTTTTGTTGTAGTAAATTCGGTTACTGTGGAACTACAGATGCGTACTGCGGTGACGGCTGTCAATCAGGACCCTGTAGATCTGGTGGGGGAGGGGGTGGAGGTGGTGGTGGTGGAGGAGGCGGTAGTGGTGGTGCTAACGTAGCCAACGTGGTTACAGATGCCTTTTTCAACGGCATCAAGAATCAAGCAGGATCCGGCTGCGAAGGAAAAAATTTCTATACCCGTAGCGCCTTTTTGAGTGCAGTAAACGCGTATCCAGGATTTGCGCATGGAGGGACCGAGGTAGAAGGAAAGCGCGAAATAGCGGCTTTTTTCGCGCATGTTACACATGAGACCGGCCACTTTTGCTATATCAGCGAAATAAATAAGTCAAACGCGTACTGTGATGCATCAAACAGACAATGGCCTTGCGCAGCTGGGCAGAAATACTATGGCCGCGGCCCTTTACAGATAAGCTGGAATTATAATTATGGCCCAGCAGGGAGAGACATTGGGTTCAATGGACTGGCAGACCCTAACCGTGTAGCCCAGGATGCAGTGATTGCATTCAAGACAGCCCTGTGGTTTTGGATGAATAATGTCCACGGCGTGATGCCGCAGGGTTTCGGCGCCACTATCCGGGCCATCAACGGGGCTTTAGAATGCAACGGCAACAACCCAGCTCAAATGAACGCTCGTGTAGGGTATTATAAACAATATTGCCAGCAATTGCGTGTGGATCCAGGACCTAATTTAATCTGTTAATAAGTCGACCGGCTGCTAACAAAGCCCGAAAGGAAGCTGAGTTGGCTGCTGCCACCGCTGAGCAATAACTAGCATAACCCCTTGGGGCCTCTAAACGGGTCTTGAGGGGTTTTTTGCTGAAAGCGAGACTAAGCTTTAAACTTCGGGTCATAGCTGTTTCCTG

6x His-AppA – 1536 base pairs

GTAAAACGACGGCCAGTAGCGCTATTAAAGCTTCGAAATTAATACGACTCACTATAGGGAGACCACAACGGTTTCCCTCTAGAAATAATTTTGTTTAACTTTAAGAAGGAGATATACATATGCATCACCACCATCATCACCAGAGTGAACCCGAGCTGAAGCTGGAATCGGTCGTTATTGTTAGCCGGCATGGGGTTAGAGCGCCTACGAAGGCTACGCAACTTATGCAAGATGTCACCCCGGACGCGTGGCCCACTTGGCCTGTAAAACTGGGATGGCTGACACCCCGCGGAGGGGAATTAATAGCGTACCTTGGTCACTATCAGAGACAGAGATTGGTTGCAGATGGCTTATTAGCAAAGAAAGGATGTCCACAAAGCGGACAGGTAGCTATCATTGCCGACGTTGATGAGAGAACGAGAAAGACAGGTGAGGCCTTCGCGGCAGGGTTAGCACCGGACTGCGCTATTACCGTTCACACCCAGGCTGATACGTCCAGTCCAGATCCGCTGTTCAATCCATTAAAGACCGGAGTCTGTCAGTTAGACAACGCGAACGTGACTGACGCTATACTGTCCCGTGCTGGAGGGTCTATTGCAGACTTTACAGGTCACCGGCAAACTGCATTTAGAGAGCTTGAGCGGGTCCTTAATTTCCCCCAGTCGAATTTATGTCTGAAGCGTGAGAAGCAAGACGAAAGTTGTTCGTTGACTCAAGCTCTGCCAAGTGAGTTGAAGGTGTCGGCGGATAACGTCAGTTTAACTGGGGCAGTAAGTTTGGCTAGTATGTTAACTGAAATCTTTTTACTTCAACAGGCTCAAGGAATGCCTGAACCAGGCTGGGGGCGGATCACGGATTCTCACCAGTGGAACACGTTGCTTTCGCTTCACAATGCACAATTTTATCTTTTGCAGCGGACTCCCGAGGTAGCGCGGTCTCGTGCCACCCCCTTATTAGATTTGATAAAGACGGCTCTTACACCTCACCCGCCCCAGAAACAAGCATATGGCGTCACCTTACCTACAAGCGTTTTGTTTATCGCGGGTCACGACACTAACCTTGCTAACTTAGGAGGGGCGTTAGAACTGAACTGGACATTACCTGGTCAGCCCGATAATACGCCCCCTGGAGGGGAATTGGTTTTCGAAAGATGGCGTCGTTTGTCGGATAACAGTCAATGGATACAGGTGAGTCTGGTGTTCCAGACTTTGCAGCAGATGAGAGACAAAACCCCACTTAGCCTTAACACCCCGCCGGGGGAAGTGAAACTTACTCTGGCCGGGTGTGAGGAACGTAACGCGCAAGGGATGTGTTCCCTGGCGGGTTTTACACAGATAGTTAACGAGGCTAGAATTCCTGCATGTTCACTTTAATAAGTCGACCGGCTGCTAACAAAGCCCGAAAGGAAGCTGAGTTGGCTGCTGCCACCGCTGAGCAATAACTAGCATAACCCCTTGGGGCCTCTAAACGGGTCTTGAGGGGTTTTTTGCTGAAAGCGAGACTAAGCTTTAAACTTCGGGTCATAGCTGTTTCCTG

rxYFP-linker-mCherry – 1761 base pairs

GTAAAACGACGGCCAGTAGCGCTATTAAAGCTTCGAAATTAATACGACTCACTATAGGGAGACCACAACGGTTTCCCTCTAGAAATAATTTTGTTTAACTTTAAGAAGGAGATATACATATGAGTAAGGGAGAGGAGTTATTTACGGGTGTAGTTCCGATACTTGTTGAACTGGATGGGGACGTAAATGGACATAAGTTTTCTGTATCGGGTGAAGGGGAGGGAGACGCAACCTACGGGAAATTGACGTTGAAATTTATTGTAACGACTGGTAAACTTCCCGTACCGTGGCCGACTCTTGTGACAACTTTTGCTTACGGTTTACAGTGTTTCGCCCGTTATCCAGACCATATGAAGAGACACGATTTTTTTAAGAGTGCGATGCCAGAAGGCTACGTCCAGGAACGTACAATATTCTTTAAGGACGACGGGAACTACAAAACACGCGCTGAAGTCAAGTTCGAGGGTGATACCTTAGTGAATCGTATAGAGTTGAAGGGAATAGACTTTAAGGAAGACGGTAACATTCTTGGCCACAAGTTAGAATACAACTATAACTCTCACTGTGTGTATATTGTGGCGGATAAGCAAAAGAATGGGATCAAGGTGAACTTTAAGATTCGGCACAACATCGAGGATGGAAGCGTTCAGCTGGCGGATCACTACCAGCAAAATACACCGATTGGGGATGGTCCGGTGTTGTTACCAGACAACCATTACCTTTGCTACCAATCGGCCCTTAGTAAAGATCCTAACGAAAAGCGTGACCATATGGTGCTGCTTGAGTTCGTAACGGCCGCAGGAATCACACATGGAATGCATGAGCTGTACAAGGCGGCTGCGTCACTTTCGACTGAATTGGAGTTCGGATCTGAGTTAATCCCCATATCGGTGAGTAAAGGCGAGGAGGACAATATGGCGATCATCAAAGAGTTCATGCGCTTCAAAGTCCACATGGAAGGCAGCGTTAATGGTCACGAGTTCGAAATTGAGGGCGAAGGCGAAGGTCGTCCGTATGAGGGTACACAGACCGCTAAACTGAAAGTCACGAAAGGTGGTCCACTGCCATTTGCTTGGGATATTCTGAGCCCACAGTTCATGTATGGCTCCAAAGCCTATGTGAAACATCCGGCCGATATTCCGGACTATCTGAAACTGAGCTTCCCTGAAGGGTTCAAATGGGAACGTGTGATGAACTTTGAGGATGGTGGTGTTGTGACAGTGACACAGGATTCTAGCCTGCAAGACGGTGAGTTCATCTATAAAGTGAAACTGCGTGGCACGAATTTTCCGAGTGATGGCCCGGTTATGCAGAAAAAAACGATGGGTTGGGAGGCCTCTAGTGAGCGTATGTATCCAGAAGATGGCGCTCTGAAAGGCGAAATCAAACAGCGTCTGAAACTGAAAGATGGTGGCCACTATGATGCCGAAGTGAAAACCACGTATAAAGCCAAAAAACCTGTCCAACTGCCTGGTGCCTATAACGTTAACATCAAACTGGACATCACCTCACACAATGAGGACTATACGATCGTGGAGCAGTATGAGCGTGCTGAAGGACGTCATTCTACCGGTGGTATGGATGAGCTGTATAAATAATAAGTCGACCGGCTGCTAACAAAGCCCGAAAGGAAGCTGAGTTGGCTGCTGCCACCGCTGAGCAATAACTAGCATAACCCCTTGGGGCCTCTAAACGGGTCTTGAGGGGTTTTTTGCTGAAAGCGAGACTAAGCTTTAAACTTCGGGTCATAGCTGTTTCCTG

### Coelenterazine Analogues

Structures drawn with ChemDraw Professional 16.0. IUPAC names from PubChem.

native: 8-benzyl-6-(4-hydroxyphenyl)-2-[(4-hydroxyphenyl)methyl]imidazo[1,2-a]pyrazin-3-ol

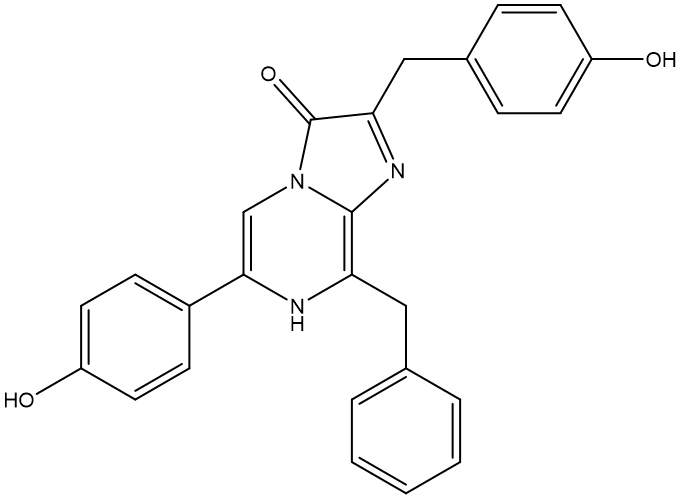

h: 2,8-dibenzyl-6-(4-hydroxyphenyl)imidazo[1,2-a]pyrazin-3-ol
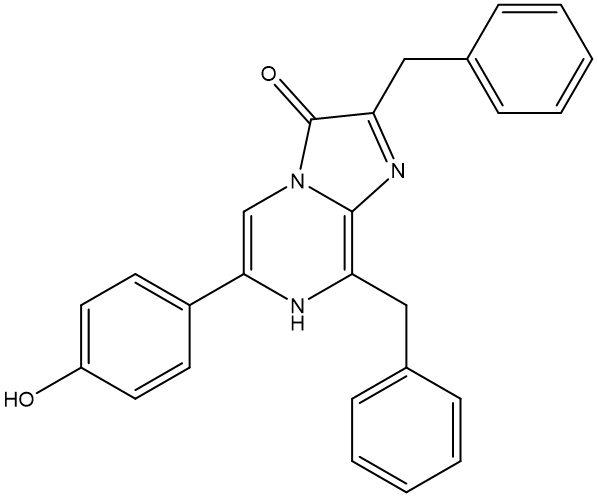

cp: 8-(cyclopentylmethyl)-6-(4-hydroxyphenyl)-2-[(4-hydroxyphenyl)methyl]imidazo[1,2-a]pyrazin-3-ol
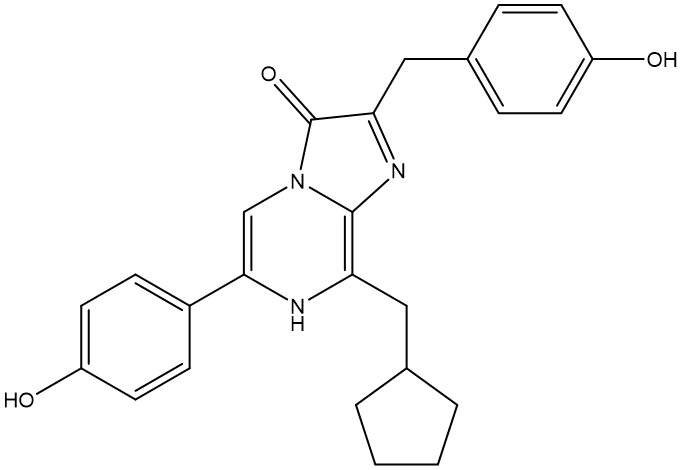

f: 8-benzyl-2-[(4-fluorophenyl)methyl]-6-(4-hydroxyphenyl)imidazo[1,2-a]pyrazin-3-ol

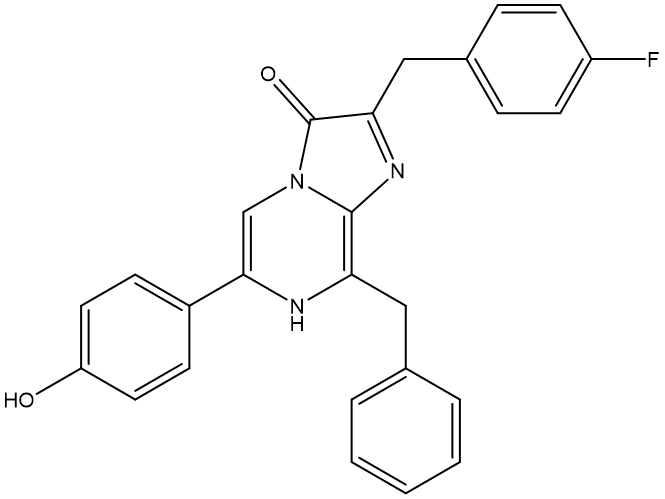

fcp: 8-(cyclopentylmethyl)-2-[(4-fluorophenyl)methyl]-6-(4-hydroxyphenyl)imidazo[1,2-a]pyrazin-3-ol

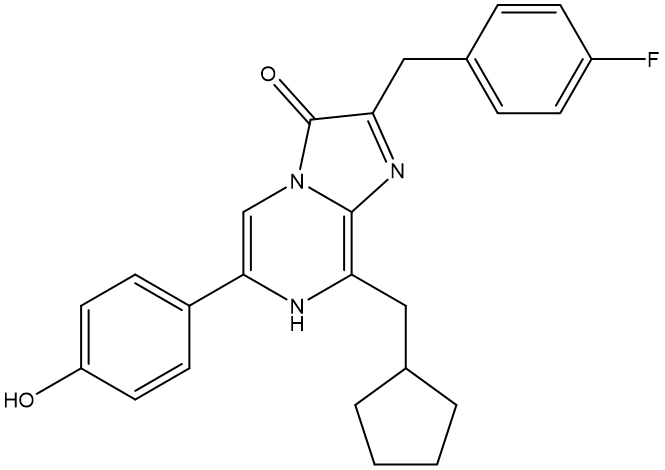

hcp: 2-benzyl-8-(cyclopentylmethyl)-6-(4-hydroxyphenyl)imidazo[1,2-a]pyrazin-3-ol

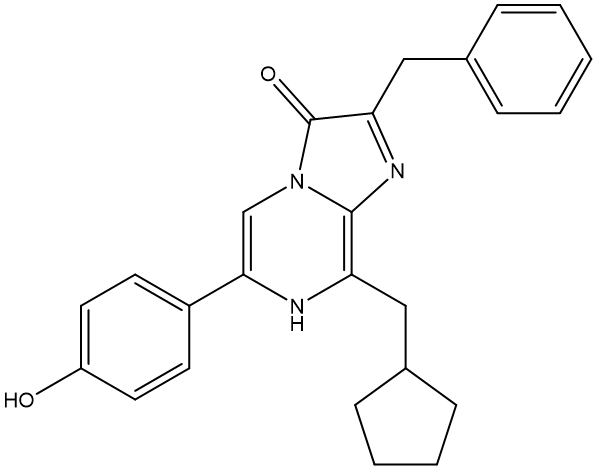

ip: 8-(isopropyl)-6-(4-hydroxyphenyl)-2-[(4-hydroxyphenyl)methyl]imidazo[1,2-a]pyrazin-3-ol

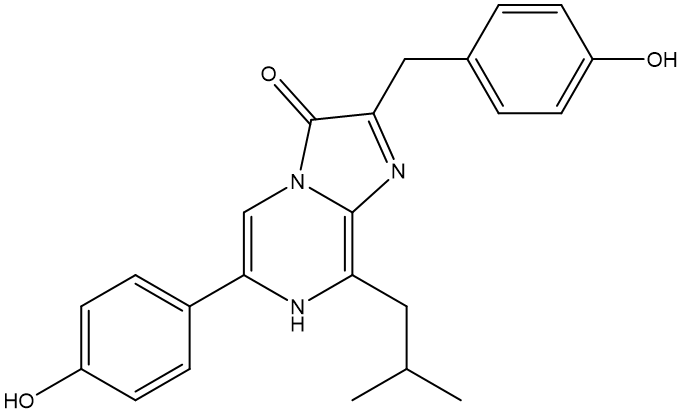

n: 8-benzyl-6-(4-hydroxyphenyl)-2-(naphthalen-1-ylmethyl)imidazo[1,2-a]pyrazin-3-ol

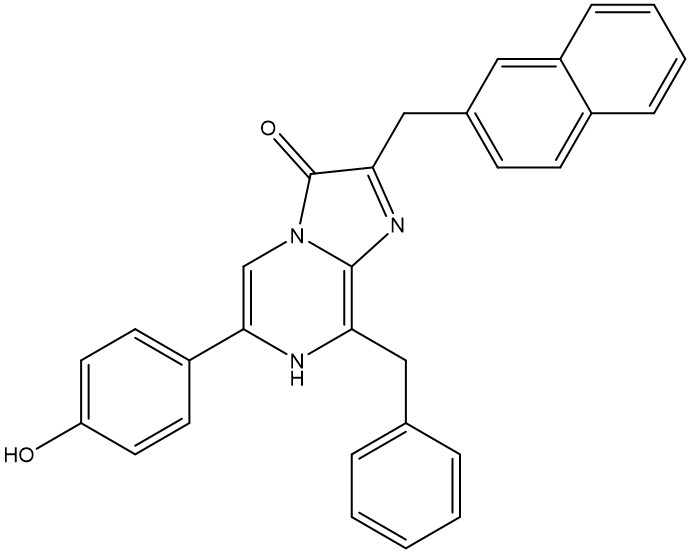

i: 8-benzyl-2-[(4-iodophenyl)methyl]-6-(4-hydroxyphenyl)imidazo[1,2-a]pyrazin-3-ol

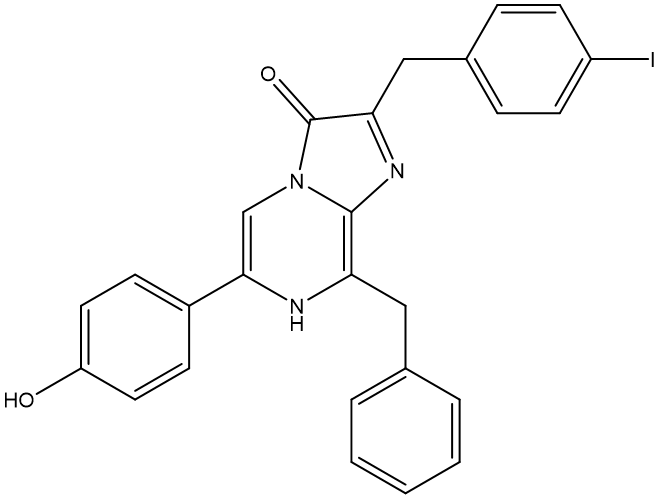
